# Cellular mechanisms of early brain overgrowth in autistic children: elevated levels of GPX4 and resistance to ferroptosis

**DOI:** 10.1101/2025.01.30.635706

**Authors:** Siwei Chen, Anna Shcherbina, Simon T. Schafer, Zoe Alexandra Mattingly, Janani Ramesh, Cyndhavi Narayanan, Sravani Banerjee, Brianna Heath, Melissa Regester, Ivette Chen, Sudhir Thakurela, Joachim Hallmayer, Ruth O’Hara, Marjorie Solomon, Christine Wu Nordahl, David G. Amaral, Sundari Chetty

**Author notes:** Correspondence to Sundari Chetty.

## Abstract

Autistic individuals with disproportionate megalencephaly (ASD-DM), characterized by enlarged brains relative to body height, have higher rates of intellectual disability and face more severe cognitive challenges than autistic children with average brain sizes. The cellular and molecular mechanisms underlying this neurophenotype remain poorly understood. To investigate these mechanisms, we generated human induced pluripotent stem cells from non-autistic typically developing children and autistic children with and without disproportionate megalencephaly. We assessed these children longitudinally from ages two to twelve years using magnetic resonance imaging and comprehensive cognitive and medical evaluations. We show that neural progenitor cells (NPCs) derived from ASD-DM children exhibit increased rates of cell survival and suppressed cell death, accompanied by heightened oxidative stress and ferrous iron accumulation. Despite these stressors, ASD-DM NPCs actively suppress apoptosis and ferroptosis by regulating proteins such as caspase-3 (CASP3), poly(ADP-ribose) polymerase 1 (PARP1), and glutathione peroxidase 4 (GPX4). Cellular ferroptotic signatures are further supported by elevated expression of selenocysteine genes, including *GPX4*, in the blood of ASD-DM children and their mothers, suggesting potential hereditary or environmental influences. Furthermore, we show that peripheral expression of *GPX4* and other selenocysteine genes correlate with cognitive outcomes (IQ). These findings underscore the role of ferroptosis in autism, pointing to potential diagnostic biomarkers and targets for intervention.

## Introduction

Autism spectrum disorder (ASD) is a highly polygenic developmental condition characterized by alterations in social communication and interaction, and restricted patterns of behaviors, interests, and activities^1,2^. The fundamental causes of autism remain unclear due to the lack of human mechanistic models and the extensive genetic and phenotypic heterogeneity associated with this condition^3^. Given this heterogeneity, identifying subtypes of autism can be helpful in understanding underlying mechanisms and identifying therapeutic targets.

Approximately 15% of autistic boys have disproportionate megalencephaly (DM) or enlarged brains relative to body height^4^. Autistic boys with DM tend to have higher rates of intellectual disability and poorer overall prognosis compared to autistic children with typical brain size^4–8^. Furthermore, prior studies have documented a strong temporal association between brain overgrowth and the onset of ASD signs^4,9–16^. These associations suggest that understanding the mechanisms regulating neurodevelopment in the ASD-DM subtype could offer potential avenues for intervening or improving outcomes.

Here, we investigate the mechanisms contributing to brain overgrowth by generating human induced pluripotent stem cells (iPSCs) from deeply phenotyped and genotyped children whom we assessed longitudinally from age 2 to 12 or beyond through the UC Davis MIND Institute Autism Phenome Project (APP). We compare non-autistic, typically developing children with brain sizes in the average range (TD-N) to two groups of autistic children: those with brain sizes in the average range (ASD-N) and those with disproportionate megalencephaly (ASD-DM) based on brain volume to height ratio greater than 1.5 standard deviations above average. We differentiated the iPSCs into neural progenitor cells (NPCs), a stage critical to neurodevelopmental alterations in autism^17^, and performed RNA-sequencing (RNA-seq) and weighted gene co-expression network analysis (WGCNA). This unbiased approach allowed us to identify differentially expressed genes and modules of highly correlated genes, thereby enhancing our understanding of the connections between gene networks and phenotypic traits.

This approach, combined with cellular phenotype analyses, revealed that NPCs derived from autistic children with DM exhibit not only increased cell proliferation – as documented in prior work^17–20^ – but also higher rates of cell survival and lower rates of cell death. NPCs from ASD-DM children display heightened oxidative stress and ferrous iron accumulation, conditions typically precipitating cell death. Despite this increased cell stress, ASD-DM NPCs actively suppress cell death pathways, including apoptosis and ferroptosis. Furthermore, ferroptotic signatures observed in iPSC-derived NPCs are also observed at the peripheral level and correlate with cognitive outcomes in autistic children.

Selenocysteine genes involved in ferroptosis and antioxidant defense mechanisms show increased expression in the blood of ASD-DM children as well as their mothers, potentially suggesting a hereditary component or shared environmental influence. Our findings uncover distinct cellular and molecular profiles in ASD-DM children and highlight potential diagnostic markers and therapeutic targets for further investigation.

## Results

### Derivation of NPCs from autistic children with divergent clinical phenotypes

To investigate the cellular mechanisms underlying brain overgrowth in autism, we isolated peripheral blood mononuclear cells (PBMCs) from twelve deeply phenotyped children across TD-N, ASD-N, and ASD-DM subgroups as part of the UC Davis MIND Institute Autism Phenome Project (APP). These children have been longitudinally assessed from ages 2 to 12 years or beyond, with comprehensive clinical, behavioral, and cognitive evaluations as well as neuroimaging^21^. The children’s development is evaluated annually for three years and periodically thereafter. Magnetic resonance imaging (MRI) at study entry between 2-4 years of age showed significantly increased brain volume relative to height (total cerebral volume, tcv; Figure 1A-C) and gray matter volume relative to height (gmv; Supplementary Figure 1A) in ASD-DM children compared to ASD-N and TD-N (p<0.01 for all comparisons). While Autism Diagnostic Observation Schedule (ADOS) calibrated severity scores were comparable between ASD-N and ASD-DM groups (Supplementary Figure 1B-D), children in the ASD-DM subgroup exhibit more severe cognitive deficits that persist with age relative to TD-N and ASD-N children. ASD-DM children showed lower gains in intellectual functioning over time as measured through intelligence quotient (IQ) scores (Figure 1D-F) and had lower adaptive behavioral skills as measured by the Vineland Adaptive Behavioral Skills assessment (Supplementary Figure 1E-H).

**Figure 1.**
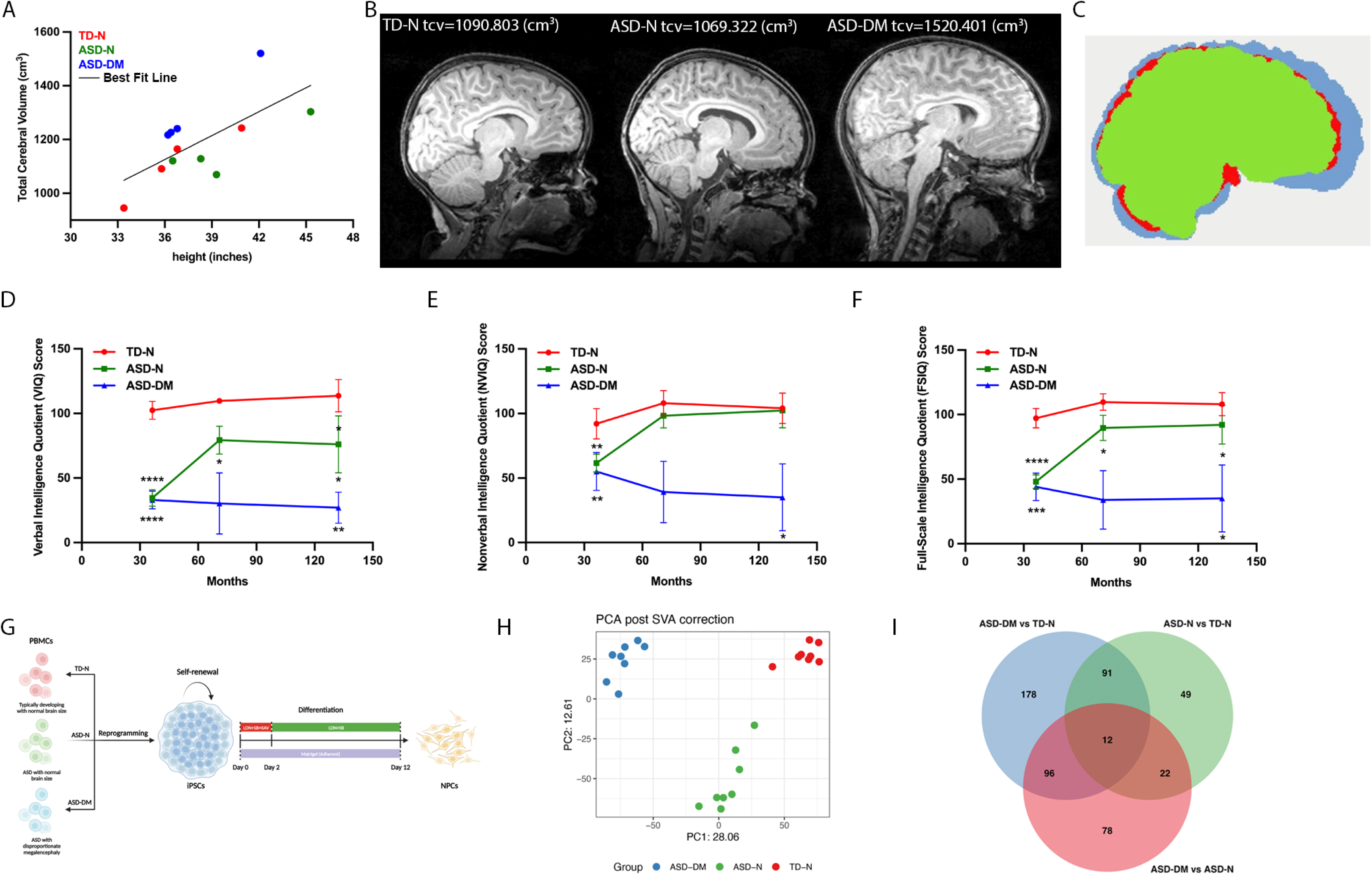
Derivation of NPSCs from autistic children with divergent clinical phenotypes. **(A)** Scatterplot of total cerebral volume (tcv) vs body height in TD-N, ASD-N and ASD-DM children. **(B)** Representative magnetic resonance imaging (MRI) of the brain from one TD-N, ASD-N, and ASD-DM participant. **(C)** Schematic representation of overlapping MRI images from each group, (TD-N, red), (ASD-N, green), and (ASD-DM, blue). **(D)** Scores of verbal intelligence quotient (IQ) in TD-N, ASD-N, and ASD-DM children with age. **(E)** Scores of non-verbal IQ in TD-N, ASD-N, and ASD-DM children with age. **(F)** Scores of full-scale IQ in TD-N, ASD-N, and ASD-DM children with age. **(G)** Schematic of neural progenitor cell (NPC) generation from induced pluripotent stem cells (iPSCs). Peripheral blood mononuclear cells (PBMCs) were isolated from TD-N, ASD-N, and ASD-DM children and reprogrammed into iPSCs. iPSCs were subsequently expanded and differentiated into NPCs using a 12-day directed differentiation protocol. **(H)** Principal component analysis (PCA) post-surrogate variable correction (SVA) of differentially expressed genes in iPSC-derived NPCs following RNA-sequencing in TD-N, ASD-N, and ASD-DM children. **(I)** Venn diagram analysis of the overlap of differentially expressed genes (FDR< 0.01, abs(log2FC)>1) across the three comparisons. All data are presented as mean ± standard deviation (*p ≤ 0.05; **p < 0.01; ***p < 0.001; **** p <0.0001). See also Figure S1, S2, and S3.

We reprogrammed the isolated PBMCs from these children into iPSCs using non-integrated Sendai viral vectors expressing the four Yamanaka factors (Oct4, Sox2, Klf4, and c-Myc) (Figure 1G, Supplementary Table 1). To investigate the mechanisms contributing to the increases in brain gray matter, we differentiated the iPSCs into neural progenitor cells (NPCs) using an established NPC differentiation protocol that gives rise to cortical neurons^22,23^. Given that prior evidence has pointed towards an important role at the neural stem cell stage in contributing to brain overgrowth in autism^17–20,24^, we focused our analyses on the NPC stage. Following 12 d of directed differentiation (Figure 1G), NPCs expressed neural markers, such as Sox1 (Supplementary Figure 2A), Pax6 (Supplementary 2B-D), and nestin, an intermediate filament protein expressed in neural stem and progenitor cells^25^ (Supplementary Figure 2E), at comparable and high levels across conditions relative to undifferentiated iPSCs. We next performed RNA-seq on the NPCs to identify genes and pathways that are differentially expressed in the autistic subpopulations relative to the TD-N individuals and potentially contributing to the DM phenotype. Following surrogate variable correction, principal component analysis (PCA) showed distinct clustering of the TD-N, ASD-N, and ASD-DM groups (Figure 1H). Differential gene expression analysis identified significantly altered genes across the three comparisons (ASD-N vs TD-N, ASD-DM vs TD-N, and ASD-DM vs ASD-N; FDR<0.01, abs(log2FC)>1) (Figure 1I, Supplementary Figure 3A-C). Notably, NPCs from ASD-DM exhibited a greater number of differentially expressed genes (DEGs) compared to TD-N (204 downregulated, 174 upregulated) relative to ASD-N (86 genes downregulated, 88 genes upregulated) (Supplementary Figure 3A-C), indicating a more pronounced transcriptomic shift in the ASD-DM neurophenotype.

Whole-genome sequencing (WGS) of the four ASD-DM children identified mutations in 12 genes linked to Gene Ontology (GO) terms associated with abnormalities in speech, social behavior, and communication, including *VPS13D*, *ABCB7*, *FIG4*, *RTTN*, *OBSCN*, *CHD8*, *PAX1*, *UROC1*, *KMT2E*, *ABCA7*, *RYR3*, and *FAN1* (Supplementary Table S2).

### Transcriptomic profiling of NPCs in ASD subtypes

We identified 103 DEGs common to ASD-DM and ASD-N compared to TD-N, while 178 genes were uniquely dysregulated in ASD-DM vs TD-N (Figure 1I). These include genes regulating mitochondrial and metabolic function, such as *NR2F2*, *PYROXD2*, *PGAM2*^26–28^. Consistent with prior studies, genes associated with autism and neurodevelopmental conditions such as *NR2F1*^29,30^ and *FZD8*, a Wnt pathway receptor implicated in brain development and size^31,32^, were upregulated in ASD-N and ASD-DM NPCs (Supplementary Figure 3A-C). Differentially expressed genes across TD-N, ASD-N, and ASD-DM NPCs revealed distinct expression profiles (Supplementary Figure 3D). Using gene set enrichment analysis (GSEA), we found downregulated genes in ASD-DM were enriched for GO terms related to translational termination, mitochondrial processes, immune pathways, ribosomal biogenesis, and rRNA metabolism (Supplementary Figure 3E). In contrast, upregulated genes in ASD conditions, particularly in ASD-DM, were associated with pathways involved in mitosis, including nuclear division, chromatid segregation, spindle assembly, as well as neural precursor cell proliferation and regulation of neuron differentiation and chromatin remodeling (Supplementary Figure 3F).

### ASD-DM NPCs exhibit increased cell viability and proliferation

To dissect the gene networks contributing to the neurophenotypes, we performed Weighted Gene Co-expression Network Analysis (WGCNA)^33,34^. This unbiased analytical approach allows for the identification of clusters (or modules) of genes that show strong co-expression, suggesting they are co-regulated and potentially involved in common biological functions or pathways. We applied WGCNA to all NPC samples from TD-N, ASD-N, and ASD-DM and found 29 distinct modules of co-expressed genes (Figure 2A-B). A significant overlap between differentially expressed genes and Module assignment were evident in eight modules (data not shown). We also quantified relationships between modules and subgroup (ASD-DM, ASD-N, and TD-N) and found several modules to be positively or negatively correlated with a particular neurophenotype (Figure 2B, values shown for Pearson correlation coefficients p<0.01). The turquoise module (Figure 2B-C) was the largest among the 29 identified (n=4295 genes), and was found to be enriched in the ASD-DM group, while showing reduction in TD-N. In line with the strong global dysregulation of cell cycle-related genes (Supplementary Figure 3), module turquoise also showed enrichment for GO terms associated with the cell cycle, cell division, cellular component organization, brain development, and chromatin organization (Figure 2D).

**Figure 2.**
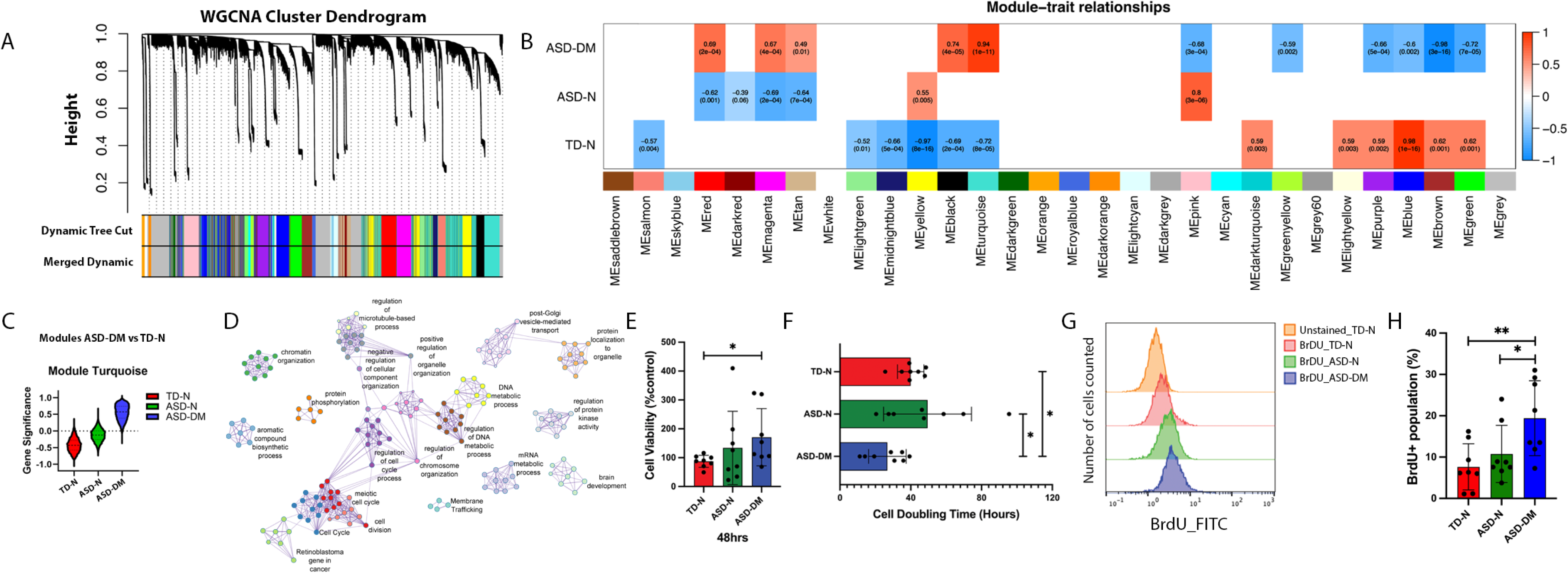
ASD-DM NPCs exhibit increased cell viability and proliferation. **(A)** WGCNA cluster dendrogram of all NPC samples, grouping genes into 29 distinct modules (row Module Colors). **(B)** Heatmap of significant overlap between DE genes and Module assignment. Relationship assignments are based on signed Pearson correlation coefficients indicating positive (red) or negative (blue) correlation of modules. Values are shown for coefficients **p ≤ 0.01. **(C)** Gene module (Module Turquoise) upregulated in ASD-DM relative to TD-N. **(D)** Module turquoise shows condition-dependent significance. GO Term Network depicts overrepresented gene sets, indicating enrichment for GO terms related to cell cycle, proliferation, and brain development. **(E)** Quantification of the percentage of viable NPCs from TD-N, ASD-N, and ASD-DM conditions measured through the MTS assay (expressed relative to the TD-N control group). **(F)** Quantification of NPC growth rates from TD-N, ASD-N, and ASD-DM conditions, assessed by cell doubling time. **(G)** Representative flow cytometric histograms of the incorporation of the thymidine analog (5-bromo-2’-deoxyuridine (BrdU) as a measure of DNA replication in TD-N, ASD-N, and ASD-DM NPCs. **(H)** Quantification of the mean fluorescence intensity of BrdU incorporation in proliferating TD-N, ASD-N, and ASD-DM NPCs. All data are presented as mean ± standard deviation (*p ≤ 0.05; **p < 0.01).

Given that several of these processes pointed towards a role for the cell cycle and cellular proliferation, we next investigated changes in cell proliferation of NPCs through various assays. The MTS assay (using the 3-(4,5-dimethylthiazol-2-yl)-5-(3-carboxymethoxyphenyl)-2-(4-sulfophenyl)-2H-tetrazolium compound) was used to measure cellular metabolic activity as an indicator of cell viability, proliferation, and cytotoxicity. We found that cellular viability was significantly increased in ASD-DM relative to TD-N NPCs (Figure 2E). We also observed a significant reduction in the cell doubling time in ASD-DM relative to TD-N and ASD-N NPCs, indicating a faster rate of proliferation or less cell death (Figure 2F).

Furthermore, incorporation of the thymidine analog 5′-bromo-2′-deoxyuridine (BrdU) demonstrated increased rates of DNA replication in ASD-DM relative to TD-N and ASD-N NPCs, as replicating cells during the S phase of the cell cycle incorporate BrdU into newly synthesized DNA (Figure 2G-H). These findings collectively corroborate the RNA-seq and WGCNA results and highlight changes in cell survival and proliferation of NPCs specifically in ASD-DM compared to ASD-N and nonautistic TD-N controls.

### ASD-DM NPCs exhibit suppressed apoptosis

WGCNA analyses also revealed the ASD-N associated module pink (n=1252 genes), which showed a strong negative correlation with ASD-DM NPCs (Figure 3A). This module was enriched for genes involved in p53 signaling, positive regulation of programmed cell death, and mitochondria-related terms (Figure 3B). Given that cell death plays an important role in regulating organ size and function through a balance of cell proliferation and programmed cell death^35^, we next investigated whether changes in cell death may contribute to the changes in brain size observed in this subgroup of autism. We further evaluated the expression patterns of important mediators of programmed cell death or apoptosis, namely cleaved PARP1 and cleaved caspase 3. Protein expression analyses through western blotting showed that ASD-DM NPCs have significantly lower levels of cleaved PARP1 and cleaved caspase 3 as compared to ASD-N and TD-N NPCs (Figure 3C-D), indicating less activation of the apoptosis pathway.

**Figure 3.**
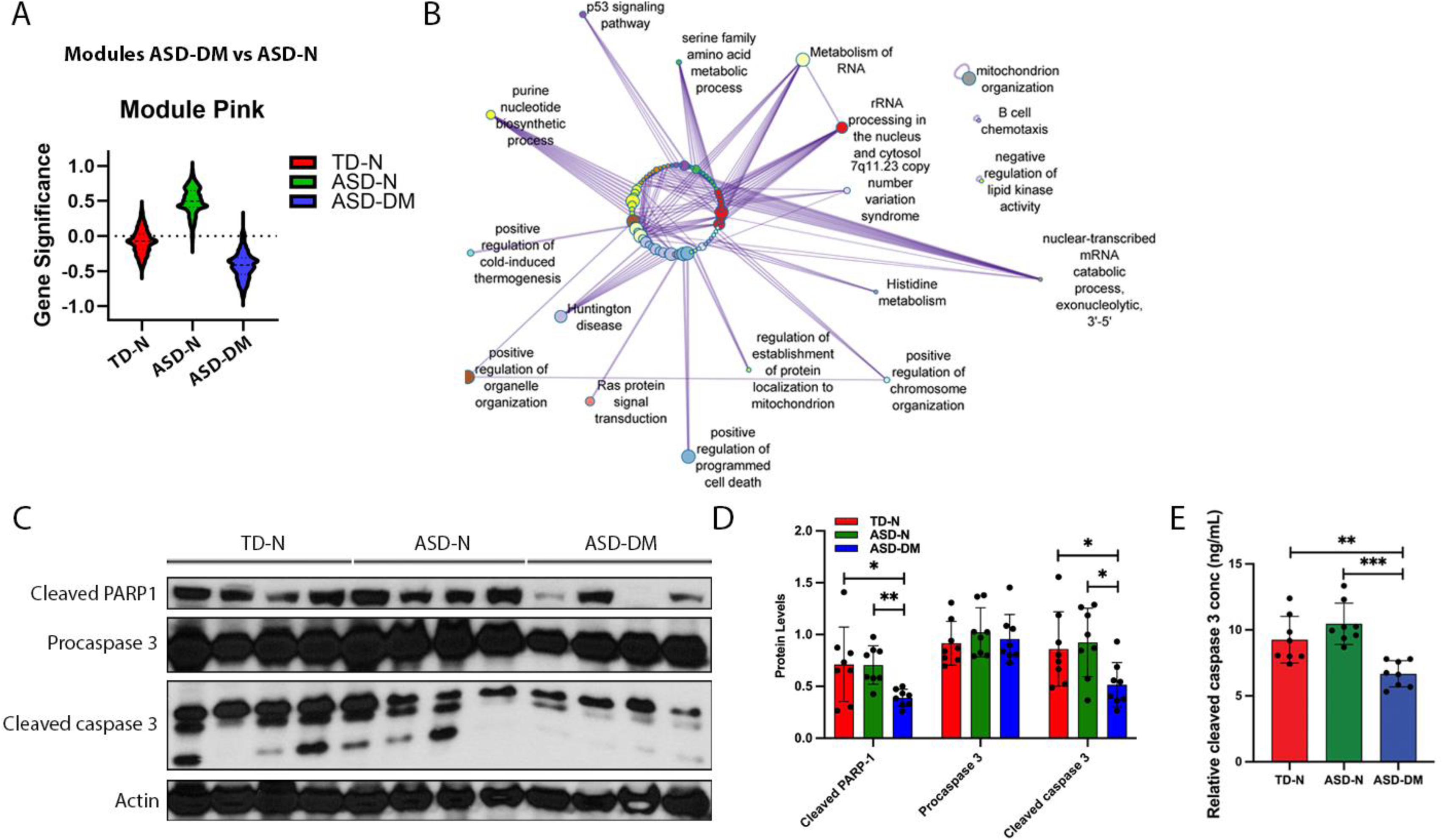
ASD-DM NPCs suppress apoptosis. **(A)** Gene module (Module Pink) downregulated in ASD-DM relative to ASD-N. **(B)** GO term network of overrepresented gene sets, highlighting enrichment for GO terms related to p53 signaling, Huntington’s disease, and positive regulation of programmed cell death. **(C)** Western blot analysis of apoptosis-related proteins in NPCs from TD-N, ASD-N, and ASD-DM children. Actin serves as a loading control. **(D)** Quantification of the expression levels of apoptosis-related proteins in NPCs from TD-N, ASD-N, and ASD-DM children, normalized to actin levels. **(E)** Expression levels of activated cleaved caspase 3 in TD-N, ASD-N, and ASD-DM NPCs as measured using an Enzyme-Linked Immunosorbent Assay (ELISA). All data are presented as mean ± standard deviation (*p ≤ 0.05; **p < 0.01; ***p < 0.001). See also Figure S4.

While procaspase 3 levels were comparable across conditions (Figure 3C-D), its cleavage and subsequent activation to caspase 3 is critical for promoting cell death. We also used an enzyme-linked immunosorbent assay (ELISA) to detect and quantify the level of cleaved caspase 3 in the NPCs, which also showed significant reductions in ASD-DM relative to TD-N and ASD-N (Figure 3E). These findings highlight lower expression levels of pro-apoptotic proteins in ASD-DM at the NPC stage. In contrast, DNA damage (measured through γ-H2AX levels; Supplementary Figure 4A-B) and proteins such as MLKL, RIP, and RIPK3 associated with another form of regulated cell death, necroptosis, were not altered (Supplementary Figure 4C-D).

### ASD-DM NPCs exhibit increased oxidative stress and suppressed ferroptosis

Ferroptosis is a regulated form of non-apoptotic cell death that plays a crucial role in various physiological and pathological contexts, including cancer, neurodegeneration, and ischemia-reperfusion injury^36–38^. Ferroptosis is distinct from other forms of cell death, such as apoptosis and necrosis, and is primarily characterized by extensive iron-dependent lipid peroxidation and an exhaustion of antioxidant defenses^36–40^. To investigate whether ferroptosis is also deregulated in ASD-DM, we measured the levels of intracellular ferrous iron (Fe^2+^) using flow cytometry in NPCs labeled with a labile Fe^2+^ dye. Strikingly, intracellular concentration of redox-active ferrous iron Fe^2+^ was significantly higher in NPCs from autistic individuals relative to those from TD-N, with the effects most pronounced in ASD-DM (Figure 4A-B).

**Figure 4.**
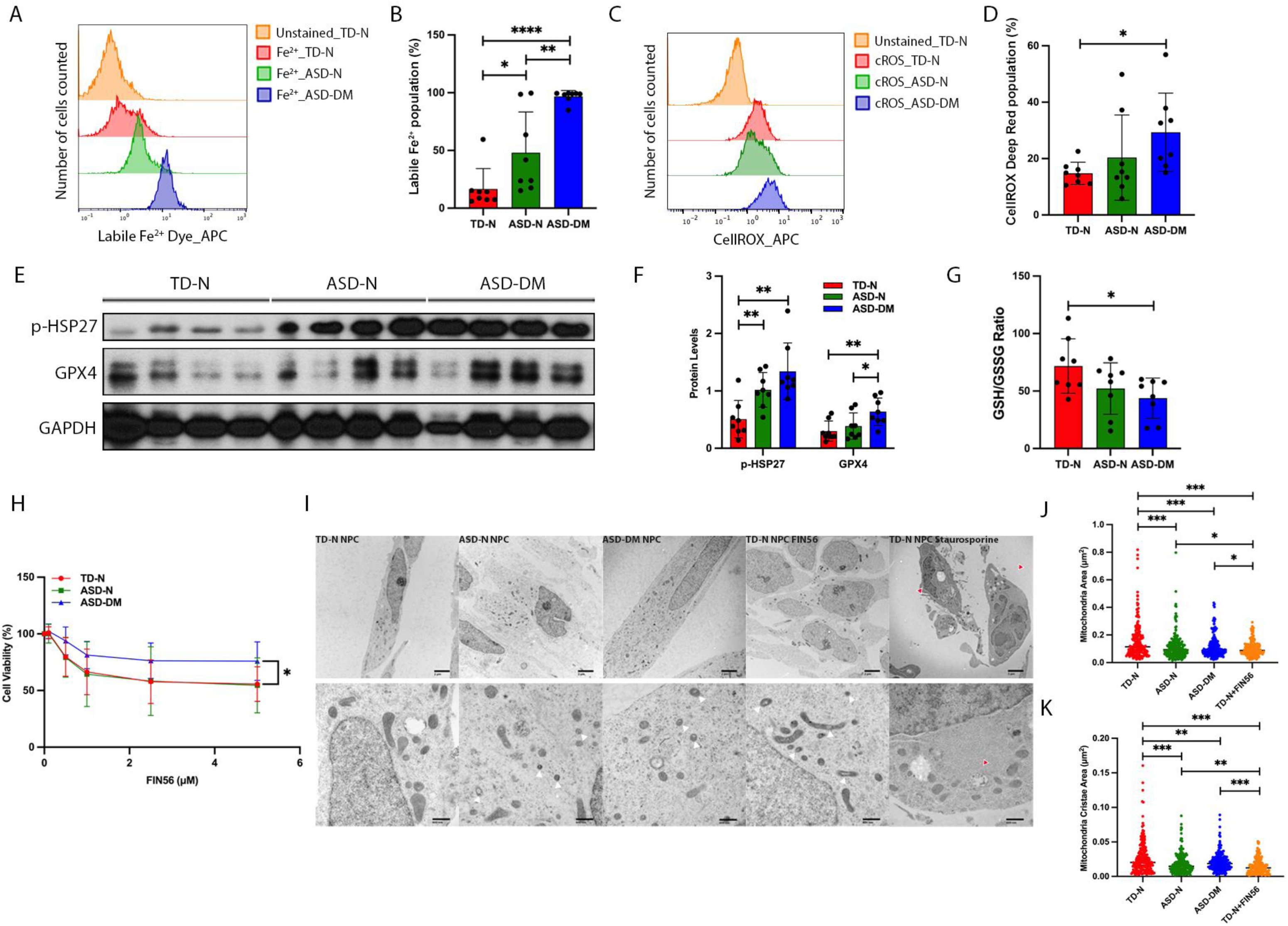
ASD-DM NPCs harbor ferroptotic signatures. **(A)** Representative flow cytometric histograms of ferrous iron (Fe^2+^) accumulation measured using a Far-red Labile Fe^2+^ Dye in TD-N, ASD-N, and ASD-DM NPCs. **(B)** Quantification of the mean fluorescence intensity of Fe^2+^ accumulation in NPCs from TD-N, ASD-N, and ASD-DM NPCs. **(C)** Representative flow cytometric histograms of cellular reactive oxygen species (cROS) measured using a CELLROX Deep Red reagent in TD-N, ASD-N, and ASD-DM NPCs. **(D)** Quantification of the mean fluorescence intensity of cROS levels in TD-N, ASD-N, and ASD-DM NPCs. **(E)** Western blot analysis of p-HSP27 and GPX4 proteins in TD-N, ASD-N, and ASD-DM NPCs. GAPDH serves as a loading control. **(F)** Quantification of protein expression levels of p-HSP27 and GPX4 in TD-N, ASD-N, and ASD-DM NPCs, normalized to GAPDH levels. **(G)** Quantification of the ratio of reduced glutathione (GSH) to oxidized glutathione (GSSG) using a Luminescent-Based Assay in TD-N, ASD-N, and ASD-DM NPCs. **(H)** Quantification of the percentage of viable NPCs from TD-N, ASD-N, and ASD-DM conditions after 48h treatment with FIN56 at increasing doses (0, 0.1, 0.5, 1, 2.5, 5 μM), measured using the MTS assay. Data are expressed relative to the corresponding untreated cell lines. **(I)** Representative electron microscopic images of TD-N, ASD-N, and ASD-DM NPCs. TD-N NPCs treated with 5 μM FIN56 for 20h were used as a positive control for ferroptosis, and the TD-N NPCs treated with 1 μM staurosporine for 3h were used as a positive control for apoptosis. Top row scale bars, 2 μm. Bottom row scale bars, 600 nm. White arrows: shrunken mitochondria. Red arrows: chromatin condensation and stress granules. **(J)** Quantification of mitochondrial area of 5 mitochondria per cell for a total of 10 cells from each sample (total of 50 mitochondria per cell line; 200 mitochondria per condition). **(K)** Quantification of mitochondrial cristae area of 5 mitochondria per cell for a total of 10 cells from each sample (total of 50 mitochondria per cell line; 200 mitochondria per condition). Individual crista areas were summed up for the total cristae surface area of each mitochondrion. **(L)** All data are presented as mean ± standard deviation (*p ≤ 0.05; **p < 0.01; ***p < 0.001; **** p <0.0001). See also Figure S5, S6, and S7.

Mitochondria play a critical role in iron metabolism, acting as key sites for both the utilization and accumulation of iron. Excessive accumulation of iron in the mitochondria can lead to oxidative stress and mitochondrial damage^41^. Using a fluorescent probe that detects ferrous iron in mitochondria, we observed an increased accumulation of Fe^2+^ in the mitochondria of NPCs from autistic individuals, particularly in ASD-DM (Supplementary Figure 5A).

Reactive oxygen species (ROS) are natural by-products of normal cellular activity and play an important role in cell signaling. However, overproduction of ROS can lead to oxidative stress and is associated with many human diseases, including cancer, cardiovascular, neurodegenerative, and metabolic disorders^42^. We next used the novel CellROX fluorogenic probe to measure ROS generation using flow cytometry and observed significantly higher levels of cellular ROS (cROS) in ASD-DM NPCs relative to TD-N (Figure 4C-D). We also used the cell permeable fluorescent and chemiluminescent probe, 2’-7’-dichlorodihydrofluorescein diacetate (DCFH-DA), to measure ROS generation and observed significantly higher levels of ROS accumulation in ASD-DM NPCs (Supplementary Figure 5B). Using a human cell stress array, we measured the relative levels of 26 different cell stress-related proteins in TD-N, ASD-N, and ASD-DM NPCs to identify differentially abundant proteins (Supplementary Figure 6A). Phosphorylated heat shock protein 27 (p-HSP27 or phoshpo-HSP27), which plays a crucial role in cellular response to stress, showed the most significant increase in ASD-DM NPCs (Supplementary Figure 6A-B). Other cell stress related proteins, such as Carbonic Anhydrase IX (CA IX), DKK4, FABP-1, NFκB, and PON3 were upregulated in ASD-DM relative to ASD-N NPCs (Supplementary Figure 6A-B). Further analysis of p-HSP27 through western blotting confirmed significantly increased expression levels in ASD-N and ASD-DM NPCs relative to TD-N NPCs (Figure 4E-F). Consistent with these findings, RNA-seq and WGCNA analyses of autistic postmortem brain samples have identified ASD-enriched modules exhibiting cortex-wide dysregulation. One upregulated module, enriched for common ASD risk variants, contains genes associated with heat-shock responses and protein folding, including HSP27^43^. In cancer, p-HSP27, as well as other cell stress related proteins such as CA IX and PON3, can facilitate stress-induced survival of cancer cells by inhibiting apoptosis and ferroptosis^44–47^.

Oxidative stress occurs when ROS production exceeds the cell’s antioxidant defenses^48^, disrupting redox balance – a risk factor for various pathologies^49^. We investigated whether this imbalance occurs in NPCs from autistic subtypes. GPX4 is a multifaceted enzyme with critical functions in cellular antioxidant defense and the regulation of cell death pathways, particularly ferroptosis. In cancer, GPX4 protects cancerous cells by preventing the accumulation of lipid hydroperoxides and thereby inhibiting ferroptosis. As a result, GPX4 may facilitate tumor growth and is a potential target for cancer therapy.

While it protects normal cells from lipid peroxidation, inhibiting GPX4 in cancer can induce ferroptosis, providing a potential therapeutic strategy^50,51^. We measured levels of GPX4 protein by western blotting and show significantly increased levels of GPX4 in ASD-DM NPCs relative to TD-N and ASD-N NPCs (Figure 4E-F). The ratio of reduced glutathione (GSH) to oxidized glutathione disulfide (GSSG) within the cell is a crucial indicator of cellular redox balance, with a high ratio of GSH to GSSG indicating a reducing environment under normal physiological conditions. However, in response to oxidative stress, this ratio shifts towards GSSG and can have significant implications for cellular health, including oxidative damage and cellular dysfunction^52^. Here, we used a specialized bioluminescent assay designed to simultaneously detect reduced (GSH) and oxidized (GSSG) forms of glutathione in the NPCs and measured the ratio of GSH to GSSG. We found the GSH:GSSG ratio to be significantly lower in ASD-DM NPCs relative to TD-N NPCs (Figure 4G), suggesting an increased level of oxidative stress in these cells. While lipid peroxidation drives cells towards ferroptosis, conditions that inhibit lipid peroxidation prevent ferroptosis^53^. Using the lipid probe 581/591 BODIPY C11, we found that lipid peroxidation levels were comparable across TD-N, ASD-N, and ASD-DM NPCs (Supplementary Figure 5C). The increased GPX4 levels in ASD-DM NPCs, along with its role in suppressing lipid peroxidation^53–56^ and the absence of elevated lipid peroxidation, suggest that ASD-DM NPCs may be protected from ferroptosis. To further investigate this, we assessed the viability of TD-N, ASD-N, and ASD-DM NPCs after treatment with the ferroptosis inducer FIN56^57^. While FIN56 significantly reduced viability in TD-N and ASD-N NPCs, ASD-DM NPCs showed increased resistance (Figure 4H).

In addition to these distinct biochemical changes, ferroptosis is characterized by distinct morphological features, such as smaller mitochondria with reduced mitochondrial cristae^58,59^. We next assessed the mitochondrial ultrastructure in NPCs using transmission electron microscopy (TEM). Autistic NPCs showed smaller mitochondria with distinct features, including reduced or absent cristae, compared to TD-N NPCs (Figure 4I-K). Treatment of TD-N NPCs with the ferroptosis inducer FIN56 led to shrunken mitochondria with diminished cristae, similar to those seen in ASD-N and ASD-DM NPCs (Figure 4I-K). These changes were not observed when TD-N NPCs were treated with staurosporine, an apoptosis inducer, highlighting a distinct response to ferroptotic conditions (Figure 4I). Cristae volume, which is positively associated with energy production^60,61^, was increased in ASD-DM NPCs compared to TD-N and ASD-N (Supplementary Figure 7A). To further explore metabolic changes, we assessed NPC activity using the Seahorse assay to measure oxygen consumption rate (OCR) and extracellular acidification rate (ECAR), indicators of mitochondrial respiration and glycolysis, respectively (Supplementary Figure 7B). ASD-DM NPCs showed significantly higher OCR, indicating increased ATP production through oxidative phosphorylation, and elevated spare respiratory capacity (SRC), which reflects a cell’s ability to enhance energy output in response to increased energy demands or stress^62^, compared to TD-N and ASD-N NPCs (Supplementary Figure 7B-C). Additionally, ECAR levels were higher in ASD-DM NPCs, suggesting that these cells function under conditions of high metabolic demand (Supplementary Figure 7D). This dual increase in oxidative phosphorylation and glycolysis points to a distinct metabolic adaptation in ASD-DM NPCs, analogous to those observed in neural stem cells ^63^ or certain cancer cells^64^ to support high energy demands during growth and proliferation or resilience under stress. These findings also align with prior studies demonstrating variability in mitochondrial dysfunction that correlates with severity in ASD^65–68^.

Taken together, despite elevated intracellular Fe^2+^ levels and oxidative stress, ASD-DM NPCs upregulate protective mechanisms compared to ASD-N, including increased GPX4 levels and resistance to ferroptosis induction, indicating that the ASD-DM cells can escape the selective instruction of cell death.

### Selenocysteine genes are upregulated in whole blood of ASD-DM children and their mothers

At the molecular level, the availability of cysteine, GSH biosynthesis, and the proper functioning of GPX4 are crucial to inhibit ferroptosis^53^. Selenium is a critical component of selenocysteine-containing proteins, including glutathione peroxidases (GPXs), thioredoxin reductases (TXNRDs), and others such as selenoproteins H and K (SELH and SELK, respectively). These selenoproteins can enhance the antioxidant capacity of cells to mitigate ferroptotic damage^69,70^. Humans have 25 known selenoproteins^71,72^ and we assessed a subset of these that are commonly associated with regulating oxidative stress, cancer, or the synthesis and function of selenoproteins^71,73–75^ in TD-N, ASD-N, and ASD-DM NPCs. Using quantitative PCR, we found that all 11 selenocysteine genes assessed (spanning *GPX*s, *TXNRD*s, and others) were upregulated in ASD-DM NPCs compared to TD-N and ASD-N (Supplementary Figure 8A-D).

Peripheral expression levels of selenium and selenoproteins, such as GPX4, may serve as potential biomarkers in various disorders, such as cancer^76^, neurodegenerative diseases^77^, sepsis^78^, and stroke^79^. Additionally, in rodent models, the mRNA expression levels of genes encoding selenoproteins in blood have been found to be expressed at comparable levels to major tissues and organs, suggesting that whole blood mRNA analysis could serve as a convenient and effective strategy for assessing these markers^80^. Given the changes in selenoproteins observed in our iPSC-derived models, we next assessed these selenocysteine genes in whole blood isolated from 7 TD-N, 8 ASD-N, and 10 ASD-DM children.

Using quantitative PCR, we found that all 11 selenocysteine genes that were assessed were upregulated in whole blood of ASD-DM children (Figure 5A-D). An average fold change of approximately 400-fold was observed for *GPX*4 in ASD-DM children compared to TD-N children. While the expression levels of common blood cell markers, including *CD45*, *CD14*, *CD33*, *CD41*, *CD61*, and *CD235a* were comparable across TD-N, ASD-N, and ASD-DM children (Supplementary Figure 9A), there was a significant increase in *CD34* expression -- a marker expressed in human hematopoietic stem and progenitor cells (HSPCS) – in ASD-DM compared to TD-N and ASD-N (Supplementary Figure 9B). Notably, hematopoietic stem cells (HSCs) have been shown to be particularly susceptible to ferroptosis^81^.

**Figure 5.**
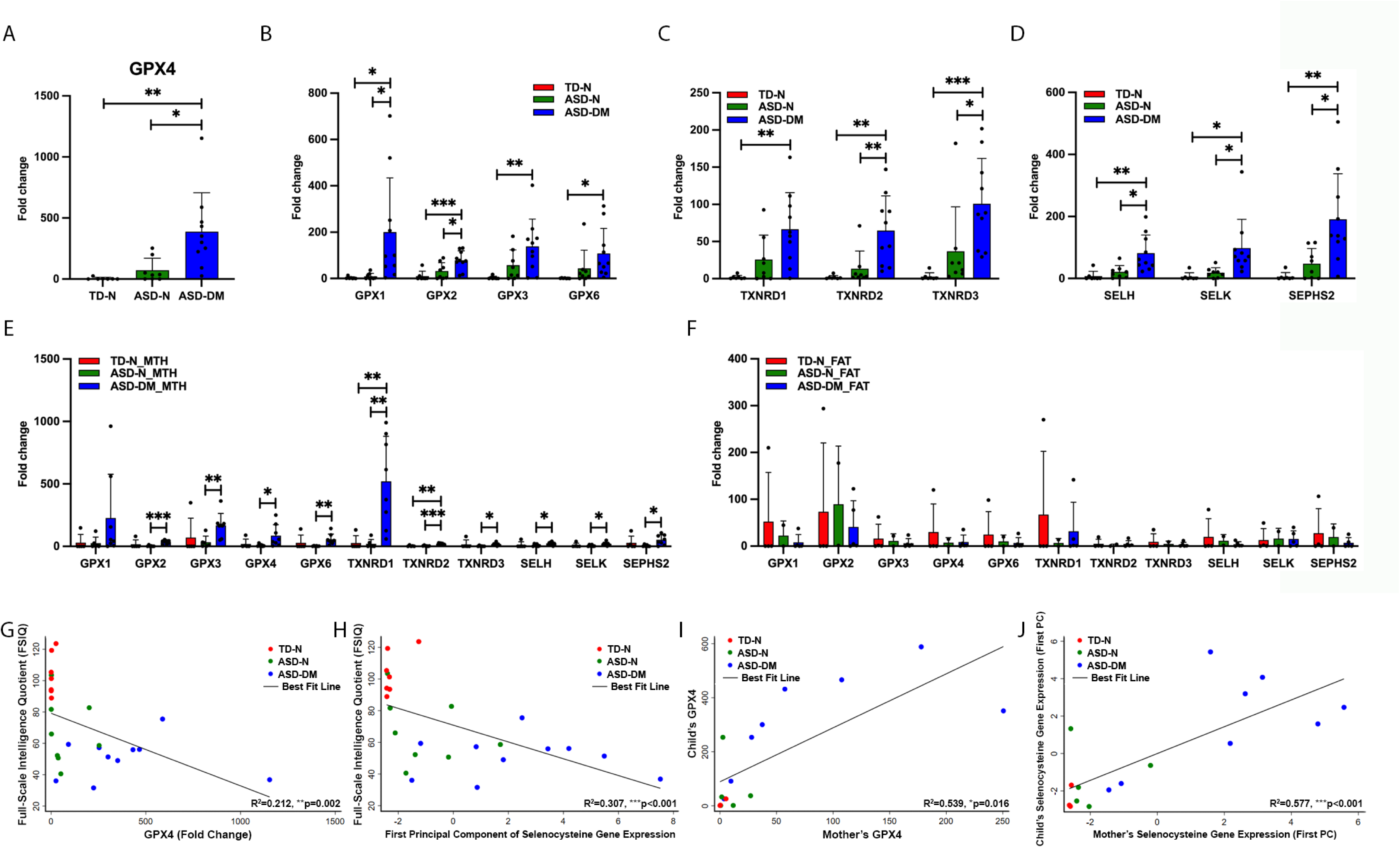
ASD-DM children and their mothers have increased expression of selenocysteine genes in their whole blood. **(A)** Quantitative real-time PCR analysis of *GPX4* gene expression in whole blood RNA from all groups (7 TD-N children, 8 ASD-N children, and 10 ASD-DM children). **(B)** Quantitative real-time PCR analysis of *GPX1*, *GPX2*, *GPX3,* and *GPX6* gene expression in whole blood RNA from all groups (7 TD-N children, 8 ASD-N children, and 10 ASD-DM children). **(C)** Quantitative real-time PCR analysis of *TXNRD1*, *TXNRD2*, and *TXNRD3* gene expression in whole blood RNA from all groups (7 TD-N children, 8 ASD-N children, and 10 ASD-DM children). **(D)** Quantitative real-time PCR analysis of *SELH*, *SELK*, and *SEPHS2* gene expression in whole blood RNA from all groups (7 TD-N children, 8 ASD-N children, and 10 ASD-DM children). **(E)** Quantitative real-time PCR analysis of expression levels of selenocysteine genes in whole blood RNA of participants’ mothers from all groups (5 TD-N mothers, 6 ASD-N mothers, and 8 ASD-DM mothers). MTH denotes mother. **(F)** Quantitative real-time PCR analysis of expression levels of selenocysteine genes in whole blood RNA of participants’ fathers from all groups (4 TD-N fathers, 2 ASD-N fathers, and 5 ASD-DM fathers). FAT denotes father. **(G)** Scatterplot of IQ vs GPX4 expression in TD-N, ASD-N, and ASD-DM children (7 TD-N children, 8 ASD-N children, and 10 ASD-DM children). **(H)** Scatterplot of IQ vs first principal component (PC) of combined selenocysteine gene expression in TD-N, ASD-N, and ASD-DM children (7 TD-N children, 8 ASD-N children, and 10 ASD-DM children). **(I)** Scatterplot of child’s GPX4 expression vs mother’s GPX4 expression in TD-N, ASD-N, and ASD-DM mother-child pairings (3 TD-N, 5 ASD-N, and 8 ASD-DM). **(J)** Scatterplot of child’s first principal component (PC) of combined selenocysteine gene expression vs mother’s first PC of combined selenocysteine gene expression in TD-N, ASD-N, and ASD-DM mother-child pairings (3 TD-N, 5 ASD-N, and 8 ASD-DM). All data are presented as mean ± standard deviation (*p ≤ 0.05; **p < 0.01; ***p < 0.001). See also Figure S8 and S9.

Given that selenocysteine gene expression can be influenced by genetic inheritance, epigenetic regulation, inflammation, and environmental or dietary factors^71,73–75^, we next investigated whether these selenocysteine genes were also upregulated in ASD-DM children’s parents. In whole blood isolated from biological mothers and fathers of TD-N, ASD-N, and ASD-DM children, we assessed the expression of these eleven selenocysteine genes and found 10 of the 11 genes to be significantly upregulated in ASD-DM mothers relative to ASD-N and/or TD-N mothers (Figure 5E), while no significant changes were observed in fathers for all eleven selenocysteine genes (Figure 5F). All blood cell markers assessed were also not significantly altered in TD-N, ASD-N, and ASD-DM mothers and fathers (Supplementary Figure 9C-D), with the exception of *CD45* which showed a modest suppression in ASD-DM fathers relative to ASD-N (Supplementary Figure 9D).

We also directly assessed the relationship between cognitive outcomes and selenocysteine gene expression. Children with higher *GPX4* mRNA levels had significantly lower IQ (Figure 5G). Notably, every child with above-median *GPX4* in the sample had an IQ below 100. We found similar results when combining all of the selenocysteine genes we analyzed using principal component analysis. Children with a higher value of the first principal component had significantly lower IQ and every child with a first principal component above the sample median had IQ below 100 (Figure 5H). These findings indicate that *GPX4* and other selenocysteine genes provide a highly sensitive diagnostic screen for cognitive disabilities.

Finally, we examined the relationship between selenocysteine gene expression in children and their mothers. We found strong positive correlations between *GPX4* levels in children and their mothers (Figure 5I). We found similarly strong correlations between the first principal component of selenocysteine gene expression between children and mothers (Figure 5J), potentially suggesting a genetic and/or environmental component to the regulation of these genes that could be familial or existing within a shared environment.

## Discussion

These findings point to certain distinct molecular and cellular mechanisms prevalent in autistic children with early brain overgrowth. The increase in iron load and GPX4 levels, shift in cellular redox homeostasis, and alterations of mitochondria in ASD-DM NPCs are all consistent with features associated with ferroptosis, a distinct form of regulated cell death that is iron-dependent. Since the initial description of ferroptosis in 2012, there have been significant advances in understanding how the metabolic state of cells influences mechanisms of cell survival and death^36,38,39,82^. Ferroptosis has been implicated in various diseases, including cancer, neurodegeneration, ischemia-reperfusion injury, and kidney failure^36–38,50,51,82^. While the precise regulation of cell survival and death is critical for proper brain development, the impact of ferroptosis on brain development and its potential contribution to neurodevelopmental conditions has yet to be explored.

Here, using iPSC-derived neural cells from children with autism, we demonstrate that NPCs in the ASD-DM neurophenotype exhibit not only increased proliferation and survival but also elevated cellular stress, including oxidative stress, ferrous iron accumulation, and mitochondrial alterations – factors typically precipitating cell death. However, these NPCs demonstrate resistance to cell death by suppressing pathways related to ferroptosis and apoptosis and elevating levels of anti-apoptotic and anti-ferroptotic proteins, including phospho-HSP27 and GPX4, similar to what is observed in cancer. This resistance is accompanied by a shift in metabolic activity, potentially to support the increased energy demand associated with increased cell survival and proliferation, resembling adaptive mechanisms observed in other stem cells^63^ and cancer cells^64^. In cancer, tumorigenic cells often outcompete normal cells through clonal expansion^83^. Future studies exploring whether specific subpopulations of NPCs in ASD-DM are particularly proliferative yet resistant to ferroptosis and/or apoptosis will be informative, as understanding these mechanisms could shed light on early neurodevelopmental processes and pathways for intervention.

Supporting these cellular findings, analyses of whole blood samples from these children and their mothers show that selenocysteine-related genes and proteins, such as GPX4, play an important role in the ASD-DM neurophenotype and could potentially serve as biomarkers, similar to those recently identified for ischemic stroke^79^. These findings demonstrate that the stem cell models developed here can mirror the molecular alterations observed in patient-derived blood samples, underscoring their utility in understanding ASD pathophysiology and identifying potential targets for intervention.

Our findings suggest that further understanding and promoting ferroptosis could have benefits in regulating cell death of unhealthy NPCs in the megalencephalic form of autism. Excessive levels of ferrous iron and iron overload contribute to cellular damage and ROS generation, leading to oxidative stress^41,84,85^. Notably, *ABCB7*, a gene involved in iron-sulfur cluster export from mitochondria, is among the genes found to have mutations in the ASD-DM group based on WGS data (Supplementary Table S2). In cancer, tumors often manipulate iron metabolism and antioxidant defense systems for their growth and survival, making them particularly vulnerable to ferroptosis induction. Exploiting this vulnerability, especially in cases where cancer cells have developed resistance to apoptosis, has proven beneficial^37,50,51,82,86^. It would be valuable to study whether compounds that have been identified to induce ferroptosis in cancer cells are helpful in eliminating unhealthy cells in ASD and related neurodevelopmental disorders. With the present study, these findings collectively suggest that regulating ferroptosis across different subtypes of autism, where there is excessive or insufficient cell loss, may be beneficial.

There is growing suggestive evidence that oxidative stress and redox imbalance play a role in the etiology of autism, but this work has primarily focused on assessments in peripheral tissues such as blood plasma and urine and has studied a heterogeneous population of autistic individuals^87–94^. These studies indicate a vulnerability to oxidative stress in autism due to low levels of plasma glutathione and reduced glutathione reserve capacity^67,89,92,95^. However, the impact of these biochemical changes on brain cells and subsequent cognitive outcomes is not well understood. By combining tools from stem cell biology with neuroimaging and clinical measures in individuals, our study extends these findings to the neural level. We demonstrate that these changes are accompanied by the accumulation of ferrous iron and particularly pronounced in children with autism characterized by brain overgrowth, yet these neural cells promote mechanisms to evade cell death. Leveraging insights from our stem cell models, we show that autistic individuals with brain overgrowth have lower IQ and higher expression of GPX4 and other selenocysteine genes. To our knowledge, this is the first study to use iPSC modeling to identify potential peripheral markers linked to cognitive outcomes.

These findings suggest that assessing selenocysteine gene expression in whole blood from larger sample sizes of autistic children and their mothers over time could help determine if these markers can lead to earlier identification of autism or predict cognitive outcomes and guide development of treatments for megalencephaly. Drawing from neurodegenerative research in Alzheimer’s and Parkinson’s diseases^96,97^, these results indicate that neuroimaging to visualize brain iron levels and distribution may warrant investigation as possible screening strategies for certain subtypes of autism. The increased risk of developing symptoms of Alzheimer’s or Parkinson’s in autistic individuals, compared to neurotypical controls^98,99^, underscores the importance of further exploring these shared mechanisms across the lifespan in autism.

## Material and Methods

### Participants

Participants were enrolled in the UC Davis MIND Institute Autism Phenome Project (APP), a longitudinal study that includes behavioral, cognitive, and neuroimaging assessments at up to 5 longitudinal time points spanning 2-19 years of age^21^. Blood specimens were acquired from participants and their biological parents. Growth measurements (height and weight) were also obtained. A subset of male children (n = 12) from the larger cohort were included based on their diagnostic status (autism or typically developing) and brain size subgroup for iPSC derivation. All aspects of the study were approved by the UC Davis Institutional Review Board and the Stem Cell Research Oversight committee. Informed consent was obtained from the guardian of each participant.

At study entry between 2-4 years of age, diagnostic confirmation for ASD was carried out using the Autism Diagnostic Interview–Revised^100^ and the Autism Diagnostic Observation Schedule (ADOS)–Generic^101^ or ADOS-2 by licensed clinical psychologists trained to research standards. The ADOS-2 provides a diagnostic cutoff and a calibrated severity score (CSS) to compare autism severity across different modules^102^. Non-autistic, non-developmentally delayed TD children (typically developing; TD) were screened using the Social Communication Questionnaire^103^ (excluded if scores were ≥11) or if they had first-degree relatives with ASD. Developmental ability was assessed using the Mullen Scales of Early Learning (MSEL)^104^. TD children were excluded if developmental scores were two or more SDs below normative means on any MSEL subscale. At study entry, ratio developmental quotients (mental age/chronological age *100) were calculated from the MSEL to provide nonverbal, verbal, and combined IQ estimates. At subsequent longitudinal visits, nonverbal, verbal, and full-scale IQ was assessed using the Differential Abilities Scales-II (DAS-II^105^).

### Neuroimaging

Magnetic resonance imaging (MRI) scans were acquired during natural nocturnal sleep at the UC Davis Imaging Research Center on a 3T Siemens TIM Trio whole-body MRI system using an 8-channel head coil. A 3-dimensional T1-weighted magnetization-prepared rapid acquisition gradient-echo (MPRAGE) sequence (TR 2,170 milliseconds; TE 4.86 milliseconds; matrix 256 × 256; 192 slices in the sagittal direction; 1.0-mm isotropic voxels) was acquired. All images were corrected for hardware-induced distortions using a calibration phantom scanned at the end of each MRI session, as previously described^106^.

### Brain size subgroup classification

Participants were subgrouped based on total cerebral volume (tcv) to height ratio at study entry (2-4 years of age) using previously established criteria^5,8,107,108^. Participants characterized as having disproportionate megalencephaly (ASD-DM) had a ratio of tcv to height that exceeded 1.5 SD above the mean of age and sex-matched controls from the APP. Participants with tcv to height ratios below 1.5 SD were characterized as having tcv to height ratios in the average range (ASD-N, TD-N).

### Whole Genome Sequencing (WGS)

Blood collected from all 4 ASD-DM children were sequenced using the 30X coverage WGS through a collaboration with MSSNG. Raw data including FASTQ and VCF files can be accessed through the MSSNG access agreement: https://research.mss.ng Whole-genome sequencing, read mapping, and variant identification (SI Appendix, Table S2) were performed as part of the MSSNG consortium^109–111^. Reads were aligned to the Genome Reference Consortium Human Build 38 (GRCh38) reference sequence.

### PBMC isolation

PBMCs (SI Appendix, Table S1) of typically developing and autistic individuals were isolated from peripheral blood as part of the APP. In brief, PBMCs were isolated by standard density gradient centrifugation with Lymphocyte Separation Medium (LSM, Corning) by centrifuging at 1700 RMP with low break at room temperature (RT) for 30 min. The “buffy coat” containing the PBMCs was then collected and rinsed and centrifuged with Hanks’ Balance Salt Solution (HBSS, Gibco). PBMCs were then resuspended and filtered through a 40 µm cell strainer. The isolated PBMCs were then counted for viability and concentration, resuspended in cryopreservation medium, and slowly frozen for storage at - 80 °C for future use.

### Whole blood collection

Whole blood samples were collected as part of the APP. Briefly, blood specimens from typically developing or autistic children (SI Appendix, Table S3) and/or their parents were collected into PAXgene tubes (IVD, Qiagen) and stored at RT for 2 hours for total lysis of the specimen. Subsequently, tubes were stored at - 20°C for 24 hours and then at -80 °C for future use.

### iPSC Reprogramming

iPSCs were reprogrammed from PBMCs of TD-N, ASD-N, and ASD-DM individuals (SI Appendix, Table S1). Briefly, 2x10^5^ PBMCs were transduced with Sendai viruses (CytoTune 2.0 kit, Thermo Fisher, USA) containing OCT3/4, SOX2, KLF4, and cMYC following the manufacturer’s protocol. Reprogrammed colonies were monitored for morphology consistent with pluripotent stem cells and picked manually for expansion. iPSC colonies were validated for the absence of Sendai virus and transgene integration through passaging the iPSCs for at least 10 passages and validated for pluripotency marker expression.

### Karyotype

Standard G-banding karyotype analyses were performed by Cell Line Genetics (Madison, WI, USA).

### Characterization and maintenance of iPSCs

All iPSC lines were cultured and maintained in mTeSR+ (Stem Cell Technologies) at 37 °C and 5% CO2 on Cultrex (R&D Systems, Bio-Techne) coated plates with 10 μM ROCK inhibitor (Y27632, Selleck Chemicals). All iPSC lines were routinely evaluated for pluripotency by assessing OCT3/4 and NANOG expression prior to differentiation.

### Regulatory and Institutional Review

All human pluripotent stem cell experiments were conducted in accordance with experimental protocols approved by the Stanford Stem Cell Research Oversight Committee and the Massachusetts General Hospital Institutional Biosafety Committee.

### Generation of NPCs

All iPSC lines were differentiated into cortical NPCs as previously described^23,24^. Briefly, the iPSCs were plated at a high-density monolayer onto Geltrex (Thermo Fisher)-precoated wells. When the cells were 90% confluent, neuroectoderm differentiation began in Essential 6 Media (Thermo Fisher) supplemented with small chemical inhibitors of TGF, SMAD, and Wnt pathways. iPSCs were treated with 500 nM LDN193189 (Tocris), 10 μM SB431542 (Tocris), and 2 μM XAV939 (Tocris) for the first three days and then 500 nM LDN193189 and 10 μM SB431542 for the remaining nine days of differentiation. Medium was freshly made and replaced daily.

### Expansion of NPCs

Following day 12 of NPC differentiation, NPCs were rinsed and dissociated using Accutase for subsequent expansion. Cells were collected and centrifuged at 1200 rpm for 10 min at RT. Subsequently, NPCs were resuspended in NPC medium, consisting of DMEM/F-12 + GlutaMAX Medium (Gibco) with B27 and N2 supplements and 20 ng/mL of FGF-2 (Gibco) and ROCK inhibitor and replated at a density of 2 million cells per well of a 6-well plate. NPCs were fed every other day with NPC medium with FGF-2 for maintenance and passaged at confluence.

### Mycoplasma Detection

iPSCs and differentiated cells were routinely confirmed to be mycoplasma-free using the Mycoplasma PCR Detection Kit (Abcam) and MycoAlert Detection Kit (Lonza Bioscience).

### Immunocytochemistry

Cells were rinsed twice in Phosphate buffered saline (PBS, Fisher Scientific), and then fixed in cold 4% paraformaldehyde (PFA, Santa Cruz Biotechnology) for 20 min at room temperature (RT), followed by three rinses in PBS. Blocking solution (5% normal donkey serum (NDS, Jackson ImmunoResearch), 0.3% Triton X-100 (Sigma-Aldrich) in PBS) was applied to cells for 1 h at RT. Subsequently, primary antibodies were diluted in the blocking buffer and incubated overnight at 4°C. Cells were then rinsed with PBS three times followed by incubation with secondary antibodies for 1 h at RT protected from light. Cells were stained with DAPI (4’,6-diamidino-2-phenylindole; 1:10,000 in PBS; Thermo Fisher) to visualize nuclei. Cells were then rinsed with PBS and imaged. The primary and secondary antibodies used and their dilution factors include: Rabbit-anti-PAX6 (1:500; BioLegend); Mouse-anti-Nestin (1:500; R&D Systems); Donkey anti-Rabbit 594 (1:500; Thermo Fisher); Donkey-anti-Mouse 488 (1:500; Thermo Fisher). All images were acquired using a Leica Fluorescent Microscope.

### RNA Isolation and Quantitative real-time PCR (qRT-PCR)

Total RNA was isolated from all iPSCs and iPSC-derived NPCs using the RNeasy Mini Kit (Qiagen) following the manufacturer’s instructions. RNA quality was measured using the Nanodrop spectrophotometer. Reverse transcription was then performed to obtain cDNA using the Reverse transcriptase system (Promega) using the same amount of RNA (25 ng/μL) from all samples.

qRT-PCR was conducted using the SYBR green system in 384-well plates. SYBR green mix (Bio-Rad), primers of target genes, and cDNA were mixed in each reaction. The Bio-Rad Real-Time PCR System was used to run the qRT-PCR experiment. The compatible Bio-Rad qRT-PCR analysis software was used for data analysis using ΔΔCT analysis method. Primers used in this study are listed in SI Appendix, Table S4.

### Western Blot

Cells were lysed using RIPA buffer (Thermo Fisher) with Protease and Phosphatase Inhibitor Cocktail (1:100 dilution; Thermo Fisher). Protein concentrations were determined by a Pierce BCA Protein Assay Kit (Thermo Fisher). Samples were separated by pre-cast SDS-PAGE (Bio-Rad) and transferred using a polyvinylidene difluoride (PVDF) membrane. The PVDF membrane was blocked with 5% Bovine Serum Albumin (BSA; Sigma-Aldrich; in PBST (0.1% Tween-20; Sigma-Aldrich)) at RT for 1 h and incubated with primary antibodies overnight at 4°C. Anti-rabbit or anti-mouse HRP conjugated antibodies were used to detect protein by chemiluminescence using ECL (Promega). Images were visualized by the CL-XPosure Film System (Thermo Fisher) and quantified with ImageJ software (National Institute of Health, USA). The primary antibodies assessed include: Rabbit-anti-PAX6 (1:2000; BioLegend); Apoptosis Western Blot Cocktail (1:250; Abcam); Rabbit-anti-phospho-HSP27 (1:1000; R&D Systems); Rabbit-anti-GPX4 (1:1000; Cell Signaling); Rabbit-anti-gamma H2A.X (1:100000; Abcam); Necroptosis antibody sampler kit (1:1000; Cell Signaling); Rabbit-anti-GAPDH (1:1000; Cell Signaling) and Rabbit-anti-β-actin (1:1000; Cell Signaling). The secondary antibodies used include: Goat-anti-Rabbit and Goat-anti-Mouse HRP (1:2000; Cell Signaling). Protein levels were quantified based on the intensity of the bands using ImageJ software, with target protein intensities normalized to the intensity of housekeeping genes (GAPDH or β-actin).

### MTS (3-(4,5-dimethylthiazol-2-yl)-5-(3-carboxymethoxyphenyl)-2-(4-sulfophenyl)-2H-tetrazolium) cell viability assay

The CellTiter 96 AQueous One Solution Cell Proliferation Assay was used for determining the number of viable cells. In brief, cells (5 x 10^3^ per well) were cultured in a 96-well plate (Corning) for 48 h at 37°C and incubated with a tetrazolium compound [3-(4,5-dimethylthiazol-2-yl)-5-(3-carboxymethoxyphenyl)-2-(4-sulfophenyl)-2H-tetrazolium, inner salt; MTS] for 90 min following manufacturer’s instructions of the CellTiter 96 AQueous One Solution Reagent (Promega). The absorbance was measured at a wavelength of 490 nm using a Synergy HTX plate reader (BioTek). Experiments were performed in triplicate and repeated at least two times. To assess cell viability following induction of ferroptosis, NPCs from TD-N, ASD-N, and ASD-DM were treated with FIN56 (a ferroptosis inducing agent) at increasing doses (0, 0.1, 0.5, 1, 2.5, 5 µM) for 48 hours prior to performing the MTS assay.

### Cell Doubling Time

Cells (2 x 10^5^ per well) were seeded in a 12-well plate and expanded for 72 h at 37 °C and 5% CO2. Total number of viable cells were assessed using Trypan Blue (Thermo Fisher) and quantified using the Countess II FL cell counter (Life Technologies).

### BrdU Labeling and Flow Cytometry Analysis

Cell proliferation was assessed using BrdU staining (Thermo Fisher) according to manufacturer’s instructions. Cells (2 x 10^5^ per well) were seeded in a 12-well plate and cultured overnight. After 48h, cells were treated with 10 μM BrdU for 2 h at 37 °C and 5% CO2. Subsequently, cells were rinsed with PBS and dissociated using Accutase (Thermo Fisher) at 400 x g for 5 min at 4 °C. Next, cells were resuspended at a density of 1 x 10^6^ in 100 μL Flow Cytometry Staining Buffer (FACS buffer; 1% NDS in PBS), followed by incubation with 1 X BrdU Staining Buffer (4X BrdU Staining Buffer was diluted in Fixation/Permeabilization Diluent) for 15 min at RT protected from light. Cells were washed with FACS buffer and incubated with 100 μL DNase I working solution (30% DNase I solution in FACS buffer) for one hour at 37 °C protected from light. Subsequently, cells were incubated with Anti-BrdU-FITC antibody (1:20 dilution) for 30 min at RT protected from light, followed by two rinses. Flow cytometry analysis was performed using the MACSQuant VYB Flow Cytometer and data were analyzed using the FlowJo software.

### Human Caspase-3 (active) Enzyme-Linked Immunosorbent Assay (ELISA)

Cleaved caspase 3 protein levels were measured using the Human Caspase-3 (active) ELISA kit (Invitrogen). Cells were lysed using Cell Extraction Buffer (Thermo Fisher) supplemented with Protease and Phosphatase Inhibitor Cocktail (1:100 dilution; Thermo Fisher), and protein concentrations were determined using the Pierce BCA Protein Assay Kit (Thermo Fisher). Protein samples and reconstituted Human Caspase-3 (active) Standard were prepared in Standard Diluent Buffer. 100 µL of each standard or sample was added to wells coated with anti-cleaved caspase 3 antibody and incubated for 2 h at RT. After rinses in 1X Wash Buffer, wells were incubated with 100 µL of Caspase-3 (active) Detection Antibody solution for 1 h at RT, followed by rinses and incubation with 100 µL of 1X Anti-Rabbit IgG HRP solution for 30 minutes at RT. Wells were then rinsed and developed with 100 µL of Stabilized Chromogen solution for 30 minutes at RT. The reaction was stopped with 100 µL of Stop Solution, and the absorbance was measured at 450 nm using a Synergy HTX plate reader (Bio Tek).

### Cellular Reactive Oxygen Species (cROS) levels

cROS levels were detected using a novel CellROX fluorogenic probe (Thermo Fisher) using flow cytometry. Cells (2 x 10^5^ per well) were seeded in a 12-well plate and incubated at 37 °C for 48h. Cells were then washed with PBS twice and incubated with CellROX Deep Red Reagent at a final concentration of 5 μM at 37 °C and 5% CO2 for 30 minutes. Next, cells were rinsed with PBS twice and dissociated using Accutase (Thermo Fisher) at 500 x g for 5 min at 4 °C, followed by fixation in 4% PFA for 15 min at RT. Cells were analyzed by flow cytometry using the MACSQuant VYB Flow Cytometer and data were analyzed using the FlowJo software.

### DCFDA cROS Assay

cROS levels were also quantitatively assessed using the cell-permeant reagent 2’3’-dichlorofluorescin diacetate (DCFDA, also known as H2DCFDA, DCFH-DA, and DCFH) in live cells (Abcam). Cells (2.5 x 10^4^ per well) were seeded in a black, clear-bottomed 96-well microplate and cultured overnight at 37 °C and 5% CO2. Cells were then washed with 1 X Buffer and incubated with diluted DCFDA solution at 37 °C for 45 min protected from light. Following incubation, cells were rinsed in 1 X Buffer and fluorescence was immediately measured using a Synergy HTX plate reader at an excitation/emission wavelength of 485/535 nm in endpoint mode.

### Lipid Peroxidation Assay

Lipid peroxidation rates were measured using the BODIPY 581/591 C11 lipid peroxidation sensor (Thermo Fisher) according to manufacturer’s instructions. Briefly, cells (2 x 10^5^) were seeded in 12-well plates and incubated overnight at 37 °C. The following day, cells were incubated with BODIPY 581/591 C11 reagent at a concentration of 5 μM for 30 minutes at 37 °C. Labeled cells were rinsed and analyzed on the MACSQuant VYB Flow Cytometer and data were analyzed using the FlowJo software.

### Human Cell Stress Array

The expression profile of cell stress related proteins was assessed using a Human Cell Stress Array Kit (R&D Systems) following the manufacturer’s instructions. Cell Lysates were isolated using Lysis Buffer 6 with Protease and Phosphatase Inhibitor Cocktail (1:100 dilution; Thermo Fisher). Protein concentrations were determined using the Pierce BCA Protein Assay Kit (Thermo Fisher). Protein samples were diluted using Array Buffer 6 and 500 μL Array Buffer 4 (1.5 mL per sample). 20 μL of reconstituted Detection Antibody Cocktail was added to each diluted sample and incubated for one hour at RT. The samples were then incubated with a pre-blocked membrane in Array Buffer 6 overnight at 4 °C. Streptavidin-HRP solution (1:2000 dilution) was diluted in Array Buffer 6 and used to detect cell stress-related proteins using chemiluminescent detection reagents. The signal produced at each capture spot corresponds to the amount of protein bound. Images were acquired using the CL-XPosure Film System (Thermo Fisher) and pixel density in each spot of the array was analyzed and quantified using the ImageJ software (National Institute of Health, USA).

### GSH/GSSG-Glo Assay

The ratio of reduced Glutathione (GSH) to oxidized Glutathione (GSSG) in living cells was measured using the GSH/GSSG-Glo Kit (Promega). Cells (2 x 10^4^ per well) were cultured overnight in a white, opaque, polystyrene, flat-bottom 96-well plate (Corning) at 37 °C and 5% CO2. After rinsing twice with PBS, cells were incubated with 50 μL of Total Glutathione Lysis Reagent or Oxidized Glutathione Lysis Reagent for 5 min at RT on a plate shaker. Cells were then incubated with 50 μL of Luciferin Generation Reagent for 30 min at RT. Next, cells were incubated with 100 μL of Luciferin Detection Reagent for 15 min at RT. The resulting relative luminescent unit (RLU) was measured using a Synergy HTX plate reader.

### Labile Ferrous Iron Tracking Assay

Labile iron (II) ions (Fe^2+^) were measured using a far-red fluorescent probe (Goryo Chemical) using Flow Cytometry. Cells (2 x 10^5^ per well) were seeded in a 12-well plate and incubated for 48 h. Cells were then washed with PBS and incubated with 5 μM FerroFarRed, diluted in non-serum DMEM/F-12 with GlutaMAX medium (Gibco), for 1 h at 37 °C and 5% CO2. Cells were then rinsed with PBS and harvested using Accutase (Thermo Fisher) at 500 x g for 5 min at 4 °C, and fixed with 4% PFA for 15 min at RT protected from light. Cells were then analyzed on a MACSQuant VYB Flow Cytometer and data were analyzed using FlowJo software.

### Mitochondrial Iron Detection Assay

Mitochondrial Fe^2+^ was detected using a Mito-FerroGreen fluorescent probe (Dojindo Molecular Technologies Inc) using fluorescence microscopy following the manufacturer’s instructions. Briefly, cells (2 x 10^4^ per well) were seeded in a 96-well plate and incubated for 24 h. Next, cells were rinsed twice with Hanks’ Balanced Salt Solution (HBSS) and incubated with 5 μM Mito-FerroGreen working solution (diluted in non-serum DMEM/F-12 with GlutaMAX medium (Gibco)) at 37 °C and 5% CO2 for 30 minutes. Subsequently, cells were rinsed with HBSS twice and imaged using fluorescence microscopy (Leica).

### Seahorse Assay

Mitochondrial function was assessed using the Seahorse XF Pro M FluxPak kit (Agilent). Cells (1 x 10^4^ per well) were cultured overnight in a Seahorse XF24 tissue culture plate. Following incubation, cells were rinsed with Seahorse XF base medium and incubated with 180 μL of the base medium supplemented with 10 mM Glucose (Agilent), 2 mM Glutamine (Agilent), and 1 mM Pyruvate (Agilent) for 45 min in a 37 °C non-CO2 incubator. Next, 2.5 μM Oligomycin, 2 μM FCCP, and 0.5 μM Rotenone were added to the pre-treated Seahorse XF Sensor Cartridge. The Sensor Cartridge was then loaded into the Agilent XFe96 Seahorse Analyzer for 20 minutes to allow for compound uptake. The cell plate was subsequently assembled with the Sensor Cartridge and loaded back into the Analyzer for measurement. Data were analyzed using the Seahorse Analytics software (Agilent).

### Transmission Electron Microscopy (TEM)

The morphology of mitochondria in NPCs was assessed using TEM. Briefly, NPCs were cultured until they reached 90% confluence in 10 cm dishes. TD-N NPCs treated with 5μM FIN56 were used as a positive control for ferroptosis, and TD-N NPCs treated with 1μM staurosporine were used as a positive control for apoptosis as previously described^81^. Cells were then rinsed and fixed using a fixation buffer composed of 2.0% paraformaldehyde/2.5% glutaraldehyde in 0.1M sodium cacodylate buffer (pH 7.4) with gentle rotation during infiltration. Cells were then rinsed with 0.1M cacodylate buffer (Electron Microscopy Sciences, Hatfield, PA) and post-fixed with 1% osmium tetroxide for 1 hour at RT. Cells were next harvested by scraping and stabilized in 2% agarose in PBS. Next, agarose-embedded pellets were dehydrated through a graded ethanol series, followed by brief dehydration in 100% propylene oxide. Specimens were then infiltrated overnight in a 1:1 mix of propylene oxide:Eponate resin (Ted Pella, Redding, CA) on a gentle rocker at RT. The following day, specimens were placed into fresh 100% Eponate and then embedded in 100% Eponate in flat molds; resin was allowed to polymerize for 24 to 45 hours at 60°C. Ultra-thin (70nm) sections were cut on a Leica EM UC7 ultramicrotome using a diamond knife (Diatome U.S., Fort Washington, PA), collected onto formvar-coated grids, stained with 2% uranyl acetate and Reynold’s lead citrate and examined in a JEOL JEM 1011 transmission electron microscope at 80 kV. Images were acquired using an AMT digital camera and imaging system with proprietary image capture software (Advanced Microscopy Techniques, Danvers, MA). Electron microscopy was performed at the Massachusetts General Hospital (MGH) Microscopy Core of the Program in Membrane Biology.

### TEM Image Analysis

TEM images (at 30,000x direct magnification) were analyzed in ImageJ following previously described protocols^81,112^. Briefly, images were uploaded in TIF format and scaled using the ’Set Scale’ option in the ’Analyze’ tab (1 pixel = 1 nm for these images). Mitochondrial area and cristae structures were outlined using the ’Freehand Selections’ tool, and the corresponding regions were saved in the ROI Manager. Area measurements were obtained using the ’Measure’ function in the ROI Manager, and the data were exported for subsequent analysis.

### Whole blood RNA isolation and qRT-PCR

Whole blood RNA was isolated from TD-N, ASD-N, and ASD-DM children and/or their parents using the PAXgene Blood RNA Kit (Qiagen) following the manufacturer’s instructions. RNA quality was assessed using a Nanodrop spectrophotometer. Reverse transcription to synthesize cDNA was performed using the Reverse Transcriptase System (Promega) with equivalent RNA input (25 ng/μL) across all samples.

qRT-PCR was performed using the SYBR green system on 384-well plates. The reaction mixture included SYBR green mix (Bio-Rad), primers for the target genes, and synthesized cDNA. The Bio-Rad Real-Time PCR System was used to run the qRT-PCR experiments, and data were analyzed using the ΔΔCT method with the compatible Bio-Rad qRT-PCR analysis software. Primers used in this study are listed in SI Appendix, Table S4

### RNA-sequencing library preparation

For bulk RNA-sequencing, total RNA was extracted from NPCs using the RNeasy Plus Mini Kit (Qiagen) following the manufacturer’s instructions. RNA quality was assessed using a Bioanalyzer (Agilent Technologies). Library preparation was performed using the KAPA Stranded mRNA-Seq Kit (Roche) following manufacturer’s instructions. Briefly, poly(A) mRNA was captured using magnetic oligo-dT beads, followed by first-strand cDNA synthesis using random primers. Second-strand synthesis was performed, converting the cDNA hybrid to double-stranded cDNA, incorporating deoxyuridine triphosphate (dUTP) into the second strand to facilitate strand-specific sequencing.

Subsequent steps included A-tailing to add deoxyadenosine monophosphate (dAMP) to the 3’-ends of the double-stranded cDNA, and adapter ligation, where double-stranded DNA adapters with 3’ dTMP overhangs were ligated to the A-tailed library fragments. The strand containing dUTP was not amplified, ensuring strand specificity. Library amplification was conducted using high-fidelity, low-bias PCR to enrich for fragments containing the proper adapter sequences. The resulting libraries were analyzed using a Bioanalyzer with the High Sensitivity DNA Assay (Agilent Technologies) to confirm library size distribution and detect potential adapter dimers.

Quantified libraries were pooled in equimolar amounts and sequenced on an Illumina NextSeq platform.

### Bioinformatic Analysis

FASTQ samples were assessed for quality using the FASTQC toolkit. The reads were then trimmed to 30 bases using cutadapt (v4.6). Trimmed reads were aligned to the GRCh38 reference genome via the STAR aligner (v 2.7.5b). Transcripts were quantified via RSEM (v1.3.3).

TPM (transcripts per million) values from RSEM were filtered for input into differential expression and pathway enrichment analysis. Downstream analysis was restricted to genes with TPM >=1 in at least one of the samples analyzed, and variance in TPM > 0.01 across the set of samples. Analysis was further restricted to ENSGID’s from Gencode (v43) that were assigned the following gene types in the Gencode GTF annotation: protein_coding, rRNA, ribozyme, sRNA, IG_C_gene, IG_D_gene, IG_J_gene, IG_V_gene, Mt_rRNA, Mt_tRNA, TR_C_Genne, TR_D_gene, TR_J_gene, TR_V_gene. Limma’s voom function in R (https://bioconductor.org/packages/release/bioc/html/limma.html) was applied to transform the TPM to a log2 scale and apply quantile normalization.

PCA analysis of the voom-transformed TPM revealed a sequencing batch effect. Surrogate variable analysis (SVA) (https://bioconductor.org/packages/release/bioc/html/sva.html) was applied to the log2TPM values, and surrogate variables that did not significantly correlate with cell type (NPC) or treatment group (ASD-N, ASD-DM, TD-N) were regressed out, along with sequencing batch, via edgeR’s removeBatchEffect function.

Batch-corrected log2TPM values underwent principal component analysis (PCA). Differential expression was then performed within the NPC samples via limma using the design formula Expression ∼ TreatmentGroup (TD-N, ASD-N, ASD-DM). Contrasts were constructed from each pair of groups (ASD-DM vs ASD-N, ASD-DM vs TD-N, ASD-N vs TD-N). For each contrast, genes were determined to be differentially expressed with limma FDR < 0.01 and abs (log2fold change) >1.

Gene set enrichment analysis (GSEA) was run on the MSIGDB curated gene ontology (GO) subset (v7.4 c5) to check for enrichments within the differentially expressed genes. For a given contrast, each gene was assigned a z-score, with z-score calculated from FDR and sign of the log-fold change. The ranked genes were provided to the Python package gseapy (v1.1) prerank function, with execution parameters min_size=5, max_size=300, permutation_num=1000, weighted_score_type=0. Significant enrichments were determined by prerank FDR < 0.01 and abs(normalized effect size) >1. The rrvgo toolkit (https://www.bioconductor.org/packages/release/bioc/html/rrvgo.html) with reduceSimMatrix threshold=0.7 was used to collapse redundant GO term enrichments to their parent terms.

Weighted correlation network analysis (WGCNA) was performed on the SVA-corrected log2TPM values via the R WGCNA package (https://cran.r-project.org/web/packages/WGCNA/index.html).

### Weighted Gene Co-expression Network Analysis (WGCNA)

To construct the signed gene co-expression network, we used the WGCNA 1.51 package in R. All NPC samples were included in the network. Network construction was performed by first creating a matrix of pairwise correlations between all pairs of genes across the measured samples. The adjacency matrix was then constructed by raising the co-expression measure (0.5 + 0.5 × correlation matrix) to the power of β = 12, which is interpreted as a soft threshold of the correlation matrix^33^. Based on the resulting adjacency matrix, we next calculated the topological overlap as a robust and biologically meaningful measure of network interconnectedness^113^. Average linkage hierarchical clustering was performed to group genes with highly similar co-expression relationships. Using the dynamic hybrid tree cut algorithm^114^, we defined modules as branches resulting from the tree cutting of the hierarchical clustering tree. The expression profiles for each module were summarized by representing them as the first principal component (module eigengene), thus explaining the maximum amount of variation in the module expression levels. Module membership measures were defined for each module as the correlation between gene expression values and the module eigengene (kME). This measure was used to compute the module significances as the average gene significance measure for all genes in a given module.

### Statistical Analysis

The data are presented as mean ± SD, derived from at least two independent experiments, unless noted otherwise. Statistical analysis was conducted using the GraphPad software v.5.0 (La Jolla, CA, USA). Data were analyzed using a two-tailed unpaired Student’s *t*-test, unless noted otherwise. Significance levels are denoted as follows: \**P*-values ≤ 0.05; \*\**P*-values < 0.01; \*\*\**P*-values < 0.001, \*\*\*\**P*-values<0.0001.

## Acknowledgements.

We thank all the families participating in the Autism Phenome Project at the UC Davis MIND Institute, the staff at the iPSC core facility at Stanford University, and the Flow Cytometry and Imaging Core facilities at MGH for their assistance, Megan Dennis and her laboratory for assistance with the whole genome sequencing analyses, and Scott Dixon, Steven Hyman, David Scadden, and Jayaraj Rajagopal and their laboratories, especially Trine Ahn Kristiansen, Lingli He, and Christina Mayerhofer for their assistance and feedback on the manuscript. The MGH Microscopy Core is partially supported by an Inflammatory Bowel Disease Grant DK043351 and a Boston Area Diabetes and Endocrinology Research Center (BADERC) Award

DK135043. Funding for this work was supported by the Massachusetts General Hospital, Harvard Medical School, Stanford University School of Medicine, UC Davis MIND Institute, and an Autism Center of Excellence grant awarded by the National Institute of Child Health and Development (Grant P50 HD093079).

## Author contributions

S.Chen designed the experiments, cultured and differentiated cell lines, performed and analyzed all cellular phenotypic assays, isolated and analyzed RNA from blood samples, and edited the manuscript.

A.S. and S.T. analyzed bulk RNA-seq data. S.T.S. performed and analyzed all WGCNA data. Z.A.M. and J.R. assisted with culturing cell lines and implementation of cellular assays. C.N. and S.B. assisted with culturing cell lines, processed samples for bulk RNA-sequencing, and characterized differentiated NPCs.

B.H. and M.R. consented participants and acquired PBMCS and blood samples through the APP for iPSC reprogramming and subsequent analyses. I.C. assisted with TEM analyses of mitochondrial structure. J.H. and R.O. contributed to initial project planning. M.S. contributed to initial project planning and acquired behavioral and clinical data on participants. C.W.N. and D.G.A. recruited and characterized cohorts for the APP and guided subgrouping and sample acquisition. C.W.N. collected longitudinal neuroimaging, clinical, and behavioral data on participants. D.G.A. provided leadership in project planning and funding for the APP. S.Chetty conceived the project, provided leadership and guidance throughout all stages of the project, including designing the experiments, analyzing the data, writing the manuscript, and securing funding for the work. All authors reviewed and edited the manuscript.

## Supplementary Figures

**Table S1. Clinical characteristics of iPSC subjects.**

**Table S2. Whole genome sequencing of ASD-DM individuals from whom iPSCs were derived.**

**Table S3. Clinical characteristics of whole blood RNA subjects.**

**Table S4. Primer sequences for quantitative RT-PCR.**

**Figure S1. Neuroimaging, clinical, and behavioral measurements of TD-N, ASD-N, and ASD-DM children, related to Figure 1**.

**(A)** Scatterplot of gray matter volume (gmv) vs body height in TD-N, ASD-N, and ASD-DM children.
**(B)** Scores of Autism Diagnosis Observation Schedule (ADOS) Overall Severity in the ASD-N and ASD-DM children with age.
**(C)** Scores of ADOS Social Affect Severity in the ASD-N and ASD-DM children with age.
**(D)** Scores of ADOS Restricted and Repetitive Behavior Severity in the ASD-N and ASD-DM children with age.
**(E)** Scores of Vineland Communication in the TD-N, ASD-N, and ASD-DM children with age.
**(F)** Scores of Vineland Daily Living Skills in the TD-N, ASD-N, and ASD-DM children with age.
**(G)** Scores of Vineland Socialization in the TD-N, ASD-N, and ASD-DM children with age.
**(H)** Scores of Vineland Adaptive Behavior Scales in the TD-N, ASD-N, and ASD-DM children with age.

All data are presented as mean ± standard deviation (*p ≤ 0.05; **p < 0.01).

**Figure S2. Expression of neural markers in iPSC-derived NPCs, related to Figure 1**.

**(A)** Quantitative real-time qRT-PCR analysis of *SOX1* gene expression of iPSC-derived NPCs from TD-N, ASD-N, and ASD-DM children.
**(B)** Western blot analysis of protein levels of PAX6 in iPSC-derived NPCs from TD-N, ASD-N, and ASD-DM children. GAPDH serves as a loading control.
**(C)** Quantification of PAX6 protein levels in iPSC-derived NPCs from TD-N, ASD-N, and ASD-DM children, normalized to GAPDH levels.
**(D)** Representative immunocytochemistry (ICC) images of PAX6 protein expression in TD-N, ASD-N, and ASD-DM NPCs. Scale bars, 50 μm.
**(E)** Representative ICC images of the intermediate filament protein Nestin in TD-N, ASD-N, and ASD-DM derived NPCs. Scale bars, 50 μm.

All data are presented as mean ± standard deviation (***p < 0.001; **** p <0.0001).

**Figure S3. Transcriptomic profiling of NPCs in autistic subtypes, related to Figure 1**.

**(A)** Volcano plot of ASD-N vs TD-N gene expression levels in NPCs.
**(B)** Volcano plot of ASD-DM vs TD-N gene expression levels in NPCs.
**(C)** Volcano plot of ASD-DM vs ASD-N gene expression levels in NPCs.
**(D)** Heatmap of the most significantly differentially expressed genes across TD-N, ASD-N, and ASD-DM NPCs (ranked by FDR, thresholding for abs(log2FC)>1).
**(E)** Gene ontology (GO) analysis of terms significantly downregulated in ASD conditions (ASD-N and ASD-DM) relative to the TD-N condition (FDR < 0.01, normalized effect size < -2).
**(F)** GO analysis of terms significantly upregulated in ASD conditions (ASD-N and ASD-DM) relative to the TD-N condition (FDR<0.01, normalized effect size>2).

For volcano plots, FDR threshold set at 0.01, log2 fold change threshold set at 1. For each comparison, the genes with the highest absolute value t-statistic are labeled.

GO Term enrichments were derived via GSEA with redundant terms removed via Revigo (using SimRel uniqueness threshold < 0.5).

**Figure S4. DNA damage and necroptosis profiles are comparable across TD-N, ASD-N, and ASD-DM NPCs, related to Figure 3**.

**(A)** Western blot analysis of gamma-H2AX protein in TD-N, ASD-N, and ASD-DM NPCs. GAPDH serves as a loading control.
**(B)** Quantification of gamma-H2AX protein levels in TD-N, ASD-N, and ASD-DM NPCs, normalized to GAPDH levels.
**(C)** Western blot analysis of necroptosis-related proteins in TD-N, ASD-N, and ASD-DM NPCs. GAPDH serves as a loading control.
**(D)** Quantification of necroptosis-related protein levels in TD-N, ASD-N, and ASD-DM NPCs, normalized to GAPDH levels.

All data are presented as mean ± standard deviation.

**Figure S5. Assessments of ferroptosis in autistic NPCs, related to Figure 4**.

**(A)** Representative fluorescent images of mitochondrial ferrous (Fe^2+^) iron accumulation in TD-N, ASD-N, and ASD-DM NPCs measured by the Mito-FerroGreen Dye. Scale bars, 50 μm.
**(B)** Quantification of cROS levels in TD-N, ASD-N, and ASD-DM NPCs based on the relative fluorescent intensity (RFI) of DCFDA/H2DCFDA-based cROS analysis.
**(C)** Quantification of cellular lipid peroxidation levels in TD-N, ASD-N, and ASD-DM NPCs measured by the ratio of oxidized and non-oxidized BODIPY dye.

All data are presented as mean ± standard deviation (*p ≤ 0.05).

**Figure S6. Cell stress-related proteins are up-regulated in ASD-DM NPCs, related to Figure 4**

**(A)** Representative membrane images and immunoblot analysis of 26 cell stress-related proteins in TD-N, ASD-N, and ASD-DM NPCs using a human cell stress array.
**(B)** Quantification of expression levels of cell stress-related proteins measured using a human cell stress array in TD-N, ASD-N, and ASD-DM NPCs, normalized to reference spots.

All data are presented as mean ± standard deviation (*p ≤ 0.05; **p < 0.01; ***p < 0.001).

Figure S7. ASD-DM NPCs exhibit mitochondrial adaptations and altered bioenergetics, related to Figure 4.

**(A)** Quantification of mitochondrial cristae volume, defined as the ratio of the total cristae surface area of each mitochondrion to the mitochondrial area, of 5 mitochondria per cell for a total of 10 cells from each sample (total of 50 mitochondria per cell line; 200 mitochondria per condition).
**(B)** Representative schematic of mitochondrial respiration measured by the Seahorse assay.
**(C)** Quantification of oxygen consumption rate (OCR) to assess basal respiration, ATP-linked respiration, maximal respiration, and spare respiratory capacity in TD-N, ASD-N, and ASD-DM NPCs.
**(D)** Quantification of extracellular acidification rate (ECAR) to assess glycolysis rates in TD-N, ASD-N, and ASD-DM NPCs.

All data are presented as mean ± standard deviation (*p ≤ 0.05; **p < 0.01; ***p < 0.001).

**Figure S8. ASD-DM NPCs have increased mRNA expression of selenocysteine genes, related to Figure 5**.

**(A)** Quantitative real-time PCR analysis of *GPX4* gene expression in NPCs derived from TD-N, ASD-N, and ASD-DM children.
**(B)** Quantitative real-time PCR analysis of *GPX1*, *GPX2*, *GPX3*, and *GPX6* gene expression in NPCs derived from TD-N, ASD-N, and ASD-DM children.
**(C)** Quantitative real-time PCR analysis of *TXNRD1*, *TXNRD2*, and *TXNRD3* gene expression in NPCs derived from TD-N, ASD-N, and ASD-DM children.
**(D)** Quantitative real-time PCR analysis of *SELH*, *SELK*, and *SEPHS2* gene expression in NPCs derived from TD-N, ASD-N, and ASD-DM children.

All data are presented as mean ± standard deviation (*p ≤ 0.05; **p < 0.01; ***p < 0.001).

**Figure S9. mRNA expression levels of blood cell markers in whole blood RNA samples from TD-N, ASD-N, and ASD-DM children and parents, related to Figure 5**.

**(A)** Quantitative real-time PCR analysis of gene expression of blood cell markers in whole blood RNA from all groups (7 TD-N children, 8 ASD-N children, and 10 ASD-DM children).
**(B)** Quantitative real-time PCR analysis of *CD34* gene expression in whole blood RNA from all groups (7 TD-N children, 8 ASD-N children, and 10 ASD-DM children).
**(C)** Quantitative real-time PCR analysis of gene expression of blood cell markers in whole blood RNA from participants’ mothers from all groups (5 TD-N mothers, 6 ASD-N mothers, and 8 ASD-DM mothers). MTH denotes mother.
**(D)** Quantitative real-time PCR analysis of gene expression of blood cell markers in whole blood RNA from participants’ fathers from all groups (4 TD-N fathers, 2 ASD-N fathers, and 5 ASD-DM fathers). FAT denotes father.

All data are presented as mean ± standard deviation (*p ≤ 0.05).

## References

1. Lord, C., Elsabbagh, M., Baird, G. & Veenstra-Vanderweele, J. Autism spectrum disorder. The Lancet 392, 508–520 (2018).

2. Lord, C. et al. Autism spectrum disorder. Nat Rev Dis Primers 6, 5 (2020).

3. Constantino, J. N. & Charman, T. Diagnosis of autism spectrum disorder: reconciling the syndrome, its diverse origins, and variation in expression. Lancet Neurol 15, 279–291 (2016).

4. Nordahl, C. W., et al. Brain enlargement is associated with regression in preschool-age boys with autism spectrum disorders. Proc. Natl. Acad. Sci. U.S.A. 108, 20195–20200 (2011).

5. Amaral, D. G. et al. In pursuit of neurophenotypes: The consequences of having autism and a big brain. Autism Research 10, 711–722 (2017).

6. Lainhart, J. E. et al. Head circumference and height in autism: a study by the Collaborative Program of Excellence in Autism. Am J Med Genet A 140, 2257–2274 (2006).

7. Courchesne, E. et al. Unusual brain growth patterns in early life in patients with autistic disorder: an MRI study. Neurology 57, 245–254 (2001).

8. Lee, J. K. et al. Longitudinal Evaluation of Cerebral Growth Across Childhood in Boys and Girls With Autism Spectrum Disorder. Biol Psychiatry 90, 286–294 (2021).

9. Fombonne, E., Rogé, B., Claverie, J., Courty, S. & Frémolle, J. Microcephaly and macrocephaly in autism. J Autism Dev Disord 29, 113–119 (1999).

10. Bartholomeusz, H. H., Courchesne, E. & Karns, C. M. Relationship between head circumference and brain volume in healthy normal toddlers, children, and adults. Neuropediatrics 33, 239–241 (2002).

11. Hazlett, H. C. et al. Early Brain Overgrowth in Autism Associated With an Increase in Cortical Surface Area Before Age 2 Years. Arch Gen Psychiatry 68, 467 (2011).

12. The IBIS Network et al. Early brain development in infants at high risk for autism spectrum disorder. Nature 542, 348–351 (2017).

13. Courchesne, E., Carper, R. & Akshoomoff, N. Evidence of brain overgrowth in the first year of life in autism. JAMA 290, 337–344 (2003).

14. Redcay, E. & Courchesne, E. When Is the Brain Enlarged in Autism? A Meta-Analysis of All Brain Size Reports. Biological Psychiatry 58, 1–9 (2005).

15. Qureshi, A. Y. et al. Opposing brain differences in 16p11.2 deletion and duplication carriers. J Neurosci 34, 11199–11211 (2014).

16. Courchesne, E. et al. Embryonic origin of two ASD subtypes of social symptom severity: the larger the brain cortical organoid size, the more severe the social symptoms. Molecular Autism 15, 22 (2024).

17. Schafer, S. T. et al. Pathological priming causes developmental gene network heterochronicity in autistic subject-derived neurons. Nat Neurosci 22, 243–255 (2019).

18. Mariani, J. et al. FOXG1-Dependent Dysregulation of GABA/Glutamate Neuron Differentiation in Autism Spectrum Disorders. Cell 162, 375–390 (2015).

19. Marchetto, M. C. et al. Altered proliferation and networks in neural cells derived from idiopathic autistic individuals. Mol Psychiatry 22, 820–835 (2017).

20. Wang, M. et al. Increased Neural Progenitor Proliferation in a hiPSC Model of Autism Induces Replication Stress-Associated Genome Instability. Cell Stem Cell 26, 221–233.e6 (2020).

21. Nordahl, C. W. et al. The Autism Phenome Project: Toward Identifying Clinically Meaningful Subgroups of Autism. Front. Neurosci. 15, 786220 (2022).

22. Tchieu, J. et al. A Modular Platform for Differentiation of Human PSCs into All Major Ectodermal Lineages. Cell Stem Cell 21, 399–410.e7 (2017).

23. Qi, Y. et al. Combined small-molecule inhibition accelerates the derivation of functional cortical neurons from human pluripotent stem cells. Nat Biotechnol 35, 154–163 (2017).

24. Li, J., et al. Overexpression of CD47 is associated with brain overgrowth and 16p11.2 deletion syndrome. Proc. Natl. Acad. Sci. U.S.A. l, e2005483118 (2021).

25. Elkabetz, Y. et al. Human ES cell-derived neural rosettes reveal a functionally distinct early neural stem cell stage. Genes Dev 22, 152–165 (2008).

26. Miao, W. et al. Nr2f2 Overexpression Aggravates Ferroptosis and Mitochondrial Dysfunction by Regulating the PGC-1α Signaling in Diabetes-Induced Heart Failure Mice. Mediators Inflamm 2022, 8373389 (2022).

27. Van Bergen, N. J. et al. Biallelic Variants in PYROXD2 Cause a Severe Infantile Metabolic Disorder Affecting Mitochondrial Function. IJMS 23, 986 (2022).

28. Jiang, X., Sun, Q., Li, H., Li, K. & Ren, X. The role of phosphoglycerate mutase 1 in tumor aerobic glycolysis and its potential therapeutic implications. Intl Journal of Cancer 135, 1991–1996 (2014).

29. Bertacchi, M. et al. NR2F1 regulates regional progenitor dynamics in the mouse neocortex and cortical gyrification in BBSOAS patients. The EMBO Journal 39, e104163 (2020).

30. Tsetsos, F. et al. Genome-Wide Association Study Points to Novel Locus for Gilles de la Tourette Syndrome. Biological Psychiatry 96, 114–124 (2024).

31. Chenn, A. & Walsh, C. A. Regulation of Cerebral Cortical Size by Control of Cell Cycle Exit in Neural Precursors. Science 297, 365–369 (2002).

32. Freese, J. L., Pino, D. & Pleasure, S. J. Wnt signaling in development and disease. Neurobiol Dis 38, 148–153 (2010).

33. Zhang, B. & Horvath, S. A general framework for weighted gene co-expression network analysis. Stat Appl Genet Mol Biol 4, Article17 (2005).

34. Langfelder, P. & Horvath, S. WGCNA: an R package for weighted correlation network analysis. BMC Bioinformatics 9, 559 (2008).

35. Vaux, D. L. & Korsmeyer, S. J. Cell Death in Development. Cell 96, 245–254 (1999).

36. Li, J. et al. Ferroptosis: past, present and future. Cell Death Dis 11, 88 (2020).

37. Yan, H.-F. et al. Ferroptosis: mechanisms and links with diseases. Signal Transduct Target Ther 6, 49 (2021).

38. Dixon, S. J. & Olzmann, J. A. The cell biology of ferroptosis. Nat Rev Mol Cell Biol (2024) doi:10.1038/s41580-024-00703-5.

39. Dixon, S. J. et al. Ferroptosis: An Iron-Dependent Form of Nonapoptotic Cell Death. Cell 149, 1060–1072 (2012).

40. Chen, X., Comish, P. B., Tang, D. & Kang, R. Characteristics and Biomarkers of Ferroptosis. Front. Cell Dev. Biol. 9, 637162 (2021).

41. Paul, B. T., Manz, D. H., Torti, F. M. & Torti, S. V. Mitochondria and Iron: current questions. Expert Rev Hematol 10, 65–79 (2017).

42. Snezhkina, A. V. et al. ROS Generation and Antioxidant Defense Systems in Normal and Malignant Cells. Oxidative Medicine and Cellular Longevity 2019, 1–17 (2019).

43. Gandal, M. J. et al. Broad transcriptomic dysregulation occurs across the cerebral cortex in ASD. Nature 611, 532–539 (2022).

44. Katsogiannou, M., Andrieu, C. & Rocchi, P. Heat shock protein 27 phosphorylation state is associated with cancer progression. Front Genet 5, 346 (2014).

45. Sun, X. et al. HSPB1 as a novel regulator of ferroptotic cancer cell death. Oncogene 34, 5617– 5625 (2015).

46. Schweikert, E.-M. et al. PON3 is upregulated in cancer tissues and protects against mitochondrial superoxide-mediated cell death. Cell Death Differ 19, 1549–1560 (2012).

47. Pastorekova, S. & Gillies, R. J. The role of carbonic anhydrase IX in cancer development: links to hypoxia, acidosis, and beyond. Cancer Metastasis Rev 38, 65–77 (2019).

48. Eruslanov, E. & Kusmartsev, S. Identification of ROS using oxidized DCFDA and flow-cytometry. Methods Mol Biol 594, 57–72 (2010).

49. Alfadda, A. A. & Sallam, R. M. Reactive oxygen species in health and disease. J Biomed Biotechnol 2012, 936486 (2012).

50. Yang, W. S. et al. Regulation of Ferroptotic Cancer Cell Death by GPX4. Cell 156, 317–331 (2014).

51. Zou, Y. et al. A GPX4-dependent cancer cell state underlies the clear-cell morphology and confers sensitivity to ferroptosis. Nat Commun 10, 1617 (2019).

52. Aquilano, K., Baldelli, S. & Ciriolo, M. R. Glutathione: new roles in redox signaling for an old antioxidant. Front Pharmacol 5, 196 (2014).

53. Seibt, T. M., Proneth, B. & Conrad, M. Role of GPX4 in ferroptosis and its pharmacological implication. Free Radical Biology and Medicine 133, 144–152 (2019).

54. Maiorino, M., Conrad, M. & Ursini, F. GPx4, Lipid Peroxidation, and Cell Death: Discoveries, Rediscoveries, and Open Issues. Antioxidants & Redox Signaling 29, 61–74 (2018).

55. Forcina, G. C. & Dixon, S. J. GPX4 at the Crossroads of Lipid Homeostasis and Ferroptosis. Proteomics 19, 1800311 (2019).

56. Ursini, F. & Maiorino, M. Lipid peroxidation and ferroptosis: The role of GSH and GPx4. Free Radical Biology and Medicine 152, 175–185 (2020).

57. Shimada, K. et al. Global survey of cell death mechanisms reveals metabolic regulation of ferroptosis. Nat Chem Biol 12, 497–503 (2016).

58. Gao, M. et al. Role of Mitochondria in Ferroptosis. Molecular Cell 73, 354–363.e3 (2019).

59. Wang, H., Liu, C., Zhao, Y. & Gao, G. Mitochondria regulation in ferroptosis. European Journal of Cell Biology 99, 151058 (2020).

60. Cogliati, S., Enriquez, J. A. & Scorrano, L. Mitochondrial Cristae: Where Beauty Meets Functionality. Trends in Biochemical Sciences 41, 261–273 (2016).

61. Campello, S. & Scorrano, L. Mitochondrial shape changes: orchestrating cell pathophysiology. EMBO Reports 11, 678–684 (2010).

62. Marchetti, P., Fovez, Q., Germain, N., Khamari, R. & Kluza, J. Mitochondrial spare respiratory capacity: Mechanisms, regulation, and significance in non-transformed and cancer cells. FASEB j. 34, 13106–13124 (2020).

63. Khacho, M. et al. Mitochondrial Dynamics Impacts Stem Cell Identity and Fate Decisions by Regulating a Nuclear Transcriptional Program. Cell Stem Cell 19, 232–247 (2016).

64. Vander Heiden, M. G., Cantley, L. C. & Thompson, C. B. Understanding the Warburg effect: the metabolic requirements of cell proliferation. Science 324, 1029–1033 (2009).

65. Frye, R. E. & Rossignol, D. A. Mitochondrial Dysfunction Can Connect the Diverse Medical Symptoms Associated With Autism Spectrum Disorders: Pediatric Research 69, 41R–47R (2011).

66. Rossignol, D. A. & Frye, R. E. Mitochondrial dysfunction in autism spectrum disorders: a systematic review and meta-analysis. Mol Psychiatry 17, 290–314 (2012).

67. Rossignol, D. A. & Frye, R. E. A review of research trends in physiological abnormalities in autism spectrum disorders: immune dysregulation, inflammation, oxidative stress, mitochondrial dysfunction and environmental toxicant exposures. Mol Psychiatry 17, 389–401 (2012).

68. Frye, R. E. Mitochondrial Dysfunction in Autism Spectrum Disorder: Unique Abnormalities and Targeted Treatments. Seminars in Pediatric Neurology 35, 100829 (2020).

69. Ingold, I. et al. Selenium Utilization by GPX4 Is Required to Prevent Hydroperoxide-Induced Ferroptosis. Cell 172, 409–422.e21 (2018).

70. Chen, X., Li, J., Kang, R., Klionsky, D. J. & Tang, D. Ferroptosis: machinery and regulation. Autophagy 17, 2054–2081 (2021).

71. Labunskyy, V. M., Hatfield, D. L. & Gladyshev, V. N. Selenoproteins: Molecular Pathways and Physiological Roles. Physiological Reviews 94, 739–777 (2014).

72. Shimada, B. K., Swanson, S., Toh, P. &Seale, L. A. Metabolism of Selenium, Selenocysteine, and Selenoproteins in Ferroptosis in Solid Tumor Cancers. Biomolecules 12, 1581 (2022).

73. Lu, J. & Holmgren, A. Selenoproteins. Journal of Biological Chemistry 284, 723–727 (2009).

74. Mangiapane, E., Pessione, A. & Pessione, E. Selenium and Selenoproteins: An Overview on Different Biological Systems. CPPS 15, 598–607 (2014).

75. Brown, K. & Arthur, J. Selenium, selenoproteins and human health: a review. Public Health Nutr. 4, 593–599 (2001).

76. Chen, C. et al. GPX4 is a potential diagnostic and therapeutic biomarker associated with diffuse large B lymphoma cell proliferation and B cell immune infiltration. Heliyon 10, e24857 (2024).

77. Chen, J. & Berry, M. J. Selenium and selenoproteins in the brain and brain diseases. Journal of Neurochemistry 86, 1–12 (2003).

78. Qu, G. et al. GPX4 is a key ferroptosis biomarker and correlated with immune cell populations and immune checkpoints in childhood sepsis. Sci Rep 13, 11358 (2023).

79. Zhang, L.-M. et al. Analysis and identification of oxidative stress-ferroptosis related biomarkers in ischemic stroke. Sci Rep 14, 3803 (2024).

80. Evenson, J. K., Wheeler, A. D., Blake, S. M. & Sunde, R. A. Selenoprotein mRNA Is Expressed in Blood at Levels Comparable to Major Tissues in Rats. The Journal of Nutrition 134, 2640–2645 (2004).

81. Zhao, J. et al. Human hematopoietic stem cell vulnerability to ferroptosis. Cell 186, 732–747.e16 (2023).

82. Stockwell, B. R. et al. Ferroptosis: A Regulated Cell Death Nexus Linking Metabolism, Redox Biology, and Disease. Cell 171, 273–285 (2017).

83. Greaves, M. & Maley, C. C. Clonal evolution in cancer. Nature 481, 306–313 (2012).

84. Winterbourn, C. C. Toxicity of iron and hydrogen peroxide: the Fenton reaction. Toxicology Letters 82–83, 969–974 (1995).

85. Dev, S. & Babitt, J. L. Overview of iron metabolism in health and disease. Hemodial Int 21 Suppl 1, S6–S20 (2017).

86. Mou, Y. et al. Ferroptosis, a new form of cell death: opportunities and challenges in cancer. J Hematol Oncol 12, 34 (2019).

87. Nadeem, A. et al. Dysregulated enzymatic antioxidant network in peripheral neutrophils and monocytes in children with autism. Progress in Neuro-Psychopharmacology and Biological Psychiatry 88, 352–359 (2019).

88. Eshraghi, R. S. et al. Early Disruption of the Microbiome Leading to Decreased Antioxidant Capacity and Epigenetic Changes: Implications for the Rise in Autism. Front Cell Neurosci 12, 256 (2018).

89. Rose, S. et al. Mitochondrial and redox abnormalities in autism lymphoblastoid cells: a sibling control study. FASEB J 31, 904–909 (2017).

90. Yui, K., Kawasaki, Y., Yamada, H. & Ogawa, S. Oxidative Stress and Nitric Oxide in Autism Spectrum Disorder and Other Neuropsychiatric Disorders. CNSNDDT 15, 587–596 (2016).

91. Frye, R. E. & James, S. J. Metabolic Pathology of Autism in Relation to Redox Metabolism. Biomarkers Med. 8, 321–330 (2014).

92. James, S. J. et al. Metabolic endophenotype and related genotypes are associated with oxidative stress in children with autism. American J of Med Genetics Pt B **141B**, 947–956 (2006).

93. Bjørklund, G. et al. Oxidative Stress in Autism Spectrum Disorder. Mol Neurobiol 57, 2314–2332 (2020).

94. Chauhan, A. & Chauhan, V. Oxidative stress in autism. Pathophysiology 13, 171–181 (2006).

95. Rose, S. et al. Evidence of oxidative damage and inflammation associated with low glutathione redox status in the autism brain. Transl Psychiatry 2, e134–e134 (2012).

96. Zecca, L., Youdim, M. B. H., Riederer, P., Connor, J. R. & Crichton, R. R. Iron, brain ageing and neurodegenerative disorders. Nat Rev Neurosci 5, 863–873 (2004).

97. Ward, R. J., Zucca, F. A., Duyn, J. H., Crichton, R. R. & Zecca, L. The role of iron in brain ageing and neurodegenerative disorders. The Lancet Neurology 13, 1045–1060 (2014).

98. Starkstein, S., Gellar, S., Parlier, M., Payne, L. & Piven, J. High rates of parkinsonism in adults with autism. J Neurodevelop Disord 7, 29 (2015).

99. Vivanti, G., Tao, S., Lyall, K., Robins, D. L. & Shea, L. L. The prevalence and incidence of early-onset dementia among adults with autism spectrum disorder. Autism Research 14, 2189–2199 (2021).

100. Lord, C., Rutter, M. & Le Couteur, A. Autism Diagnostic Interview-Revised: A revised version of a diagnostic interview for caregivers of individuals with possible pervasive developmental disorders. J Autism Dev Disord 24, 659–685 (1994).

101. Lord, C. et al. The autism diagnostic observation schedule-generic: a standard measure of social and communication deficits associated with the spectrum of autism. J Autism Dev Disord 30, 205–223 (2000).

102. Gotham, K., Pickles, A. & Lord, C. Standardizing ADOS scores for a measure of severity in autism spectrum disorders. J Autism Dev Disord 39, 693–705 (2009).

103. Rutter, M., Bailey, A. &Lord, C. Social communication questionnaire (SCQ). 1 manual (iv, 23 pages : illustrations ; 28 cm), 20 Current AutoScore forms, 20 Lifetime AutoScore forms (2003).

104. Mullen, E. M. & American Guidance Service. Mullen Scales of Early Learning. (AGS, Circle Pines, Minnesota, 1995).

105. Colin D. Elliott, Joseph D. Salerno, Ron Dumont, & John O. Willis. Differential Ability Scales Second Edition. (San Antonio, TX, 2007).

106. Nordahl, C. W. et al. Increased rate of amygdala growth in children aged 2 to 4 years with autism spectrum disorders: a longitudinal study. Arch Gen Psychiatry 69, 53–61 (2012).

107. Libero, L. E. et al. Persistence of megalencephaly in a subgroup of young boys with autism spectrum disorder: Megalencephaly in Boys with Autism. Autism Research 9, 1169–1182 (2016).

108. Ohta, H. et al. Increased Surface Area, but not Cortical Thickness, in a Subset of Young Boys With Autism Spectrum Disorder. Autism Res 9, 232–248 (2016).

109. Trost, B. et al. Genomic architecture of autism from comprehensive whole-genome sequence annotation. Cell 185, 4409–4427.e18 (2022).

110. C Yuen, R. K., et al. Whole genome sequencing resource identifies 18 new candidate genes for autism spectrum disorder. Nat Neurosci 20, 602–611 (2017).

111. Chan, A. J. S. et al. Genome-wide rare variant score associates with morphological subtypes of autism spectrum disorder. Nat Commun 13, 6463 (2022).

112. Lam, J. et al. A Universal Approach to Analyzing Transmission Electron Microscopy with ImageJ. Cells 10, 2177 (2021).

113. Yip, A. M. & Horvath, S. Gene network interconnectedness and the generalized topological overlap measure. BMC Bioinformatics 8, 22 (2007).

114. Langfelder, P., Zhang, B. & Horvath, S. Defining clusters from a hierarchical cluster tree: the Dynamic Tree Cut package for R. Bioinformatics 24, 719–720 (2008).

