## Supplementary figures and tables for "Cellular mechanisms of early brain overgrowth in autistic children: elevated levels of GPX4 and resistance to ferroptosis"

Figure S1

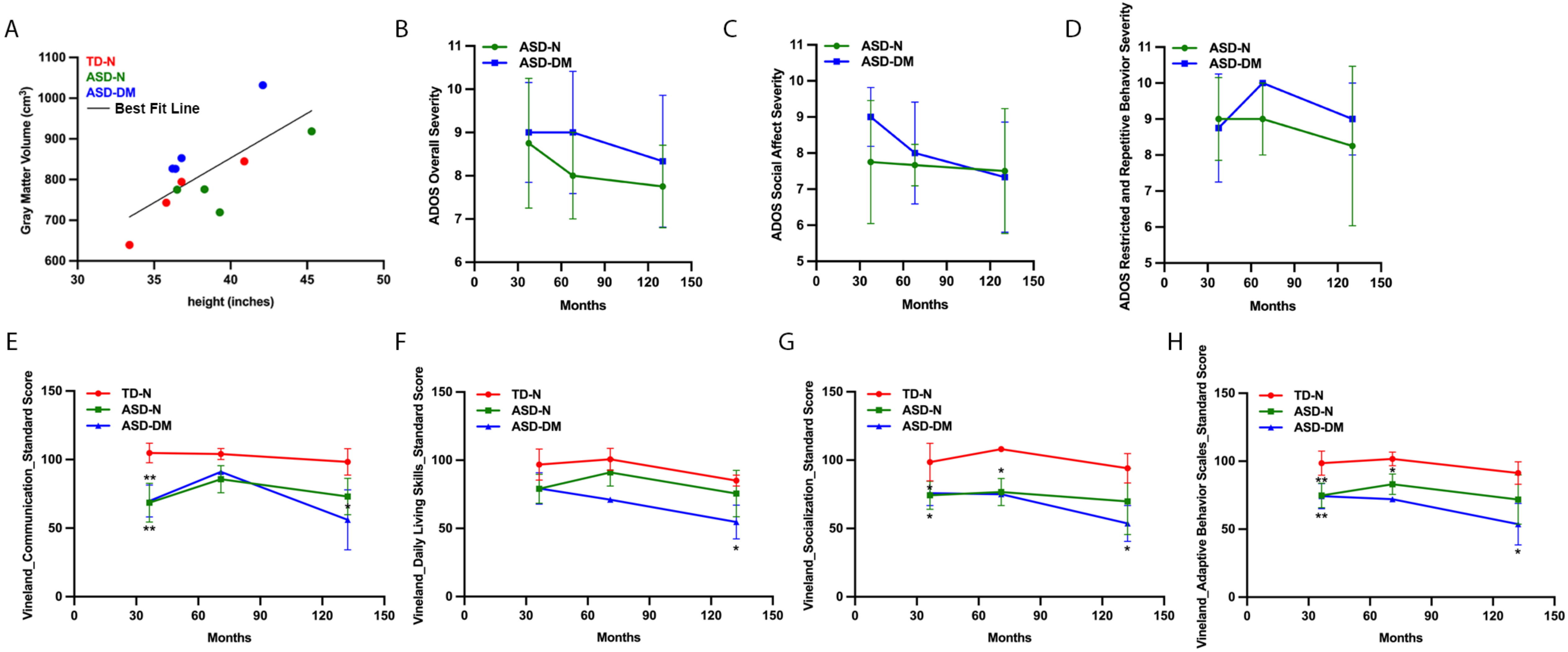

A
TD-N
ASD-N
ASD-DM
Best Fit Line
Gray Matter Volume (cm³)
height (inches)
B
ADOS Overall Severity
Months
C
ADOS Social Affect Severity
Months
D
ADOS Restricted and Repetitive Behavior Severity
Months
E
Vineland_Communication_Standard Score
Months
F
Vineland_Daily Living Skills_Standard Score
Months
G
Vineland_Socialization_Standard Score
Months
H
Vineland_Adaptive Behavior Scales_Standard Score
Months

Figure S2

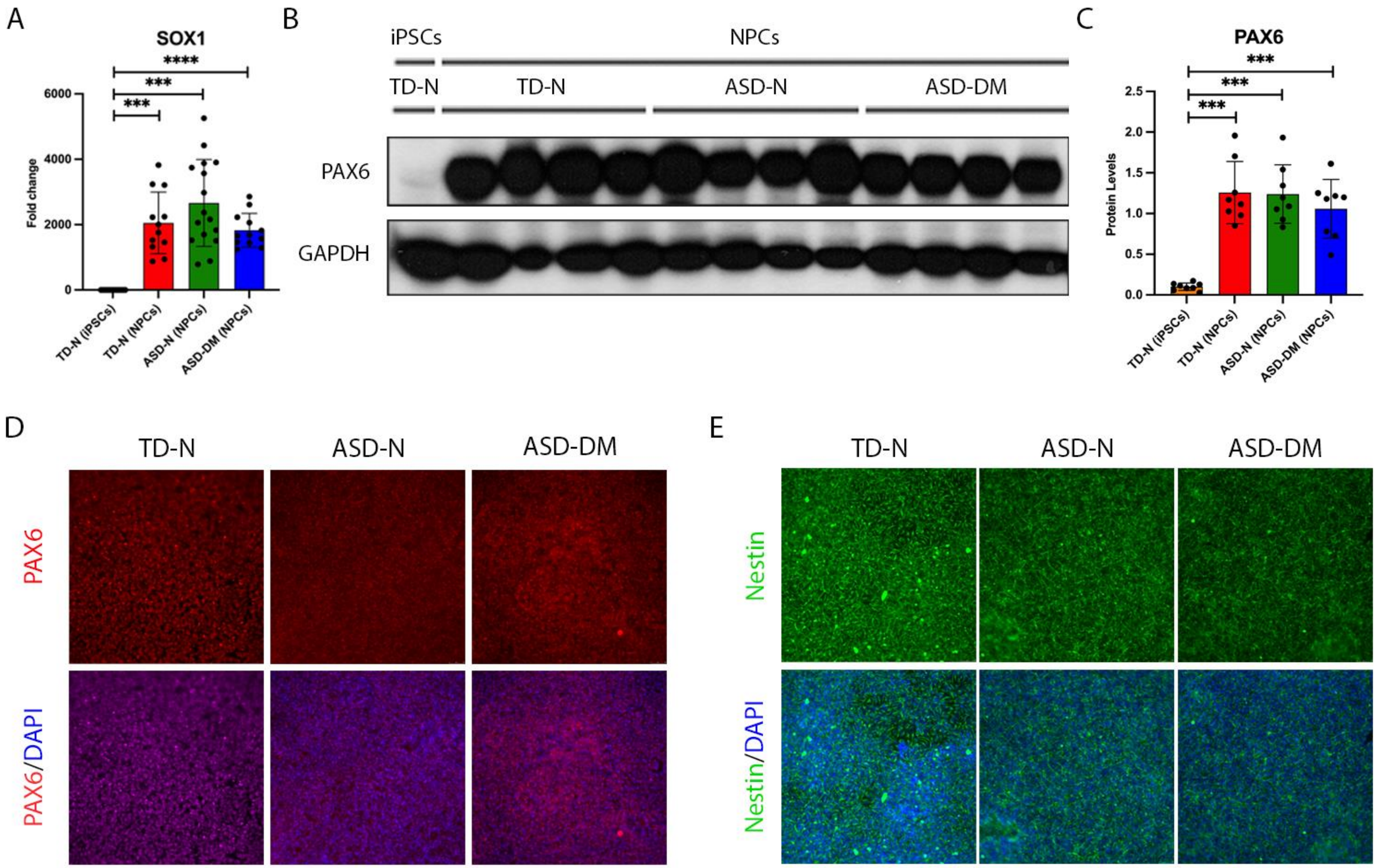

A
SOX1
Fold change
6000
4000
2000
0
****
***
***
TD-N (iPSCs)
TD-N (NPCs)
ASD-N (NPCs)
ASD-DM (NPCs)
B
iPSCs
NPCs
TD-N
TD-N
ASD-N
ASD-DM
PAX6
GAPDH
C
PAX6
Protein Levels
2.5
2.0
1.5
1.0
0.5
0.0
***
***
***
TD-N (iPSCs)
TD-N (NPCs)
ASD-N (NPCs)
ASD-DM (NPCs)
D
TD-N
ASD-N
ASD-DM
PAX6
PAX6/DAPI
E
TD-N
ASD-N
ASD-DM
Nestin
Nestin/DAPI

Figure S3

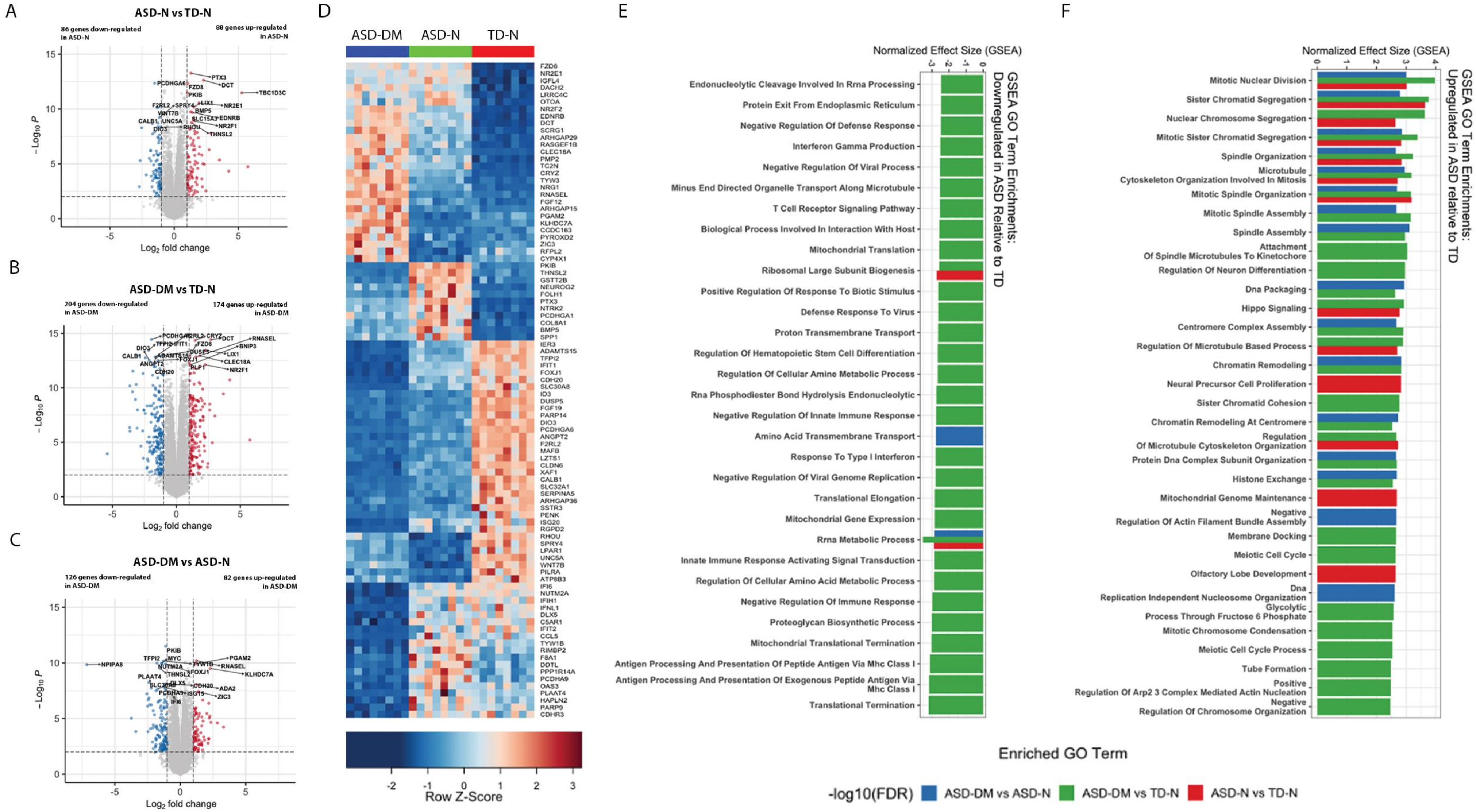

A
ASD-N vs TD-N
86 genes down-regulated in ASD-N
88 genes up-regulated in ASD-N
−Log10 P
Log2 fold change
B
ASD-DM vs TD-N
204 genes down-regulated in ASD-DM
174 genes up-regulated in ASD-DM
−Log10 P
Log2 fold change
C
ASD-DM vs ASD-N
126 genes down-regulated in ASD-DM
82 genes up-regulated in ASD-DM
−Log10 P
Log2 fold change
D
ASD-DM
ASD-N
TD-N
Row Z-Score
E
Normalized Effect Size (GSEA)
GSEA GO Term Enrichments: Downregulated in ASD Relative to TD
Endonucleolytic Cleavage Involved In Rrna Processing
Protein Exit From Endoplasmic Reticulum
Negative Regulation Of Defense Response
Interferon Gamma Production
Negative Regulation Of Viral Process
Minus End Directed Organelle Transport Along Microtubule
T Cell Receptor Signaling Pathway
Biological Process Involved In Interaction With Host
Mitochondrial Translation
Ribosomal Large Subunit Biogenesis
Positive Regulation Of Response To Biotic Stimulus
Defense Response To Virus
Proton Transmembrane Transport
Regulation Of Hematopoietic Stem Cell Differentiation
Regulation Of Cellular Amine Metabolic Process
Rna Phosphodiester Bond Hydrolysis Endonucleolytic
Negative Regulation Of Innate Immune Response
Amino Acid Transmembrane Transport
Response To Type I Interferon
Negative Regulation Of Viral Genome Replication
Translational Elongation
Mitochondrial Gene Expression
Rrna Metabolic Process
Innate Immune Response Activating Signal Transduction
Regulation Of Cellular Amino Acid Metabolic Process
Negative Regulation Of Immune Response
Proteoglycan Biosynthetic Process
Mitochondrial Translational Termination
Antigen Processing And Presentation Of Peptide Antigen Via Mhc Class I
Antigen Processing And Presentation Of Exogenous Peptide Antigen Via Mhc Class I
Translational Termination
F
Normalized Effect Size (GSEA)
GSEA GO Term Enrichments: Upregulated in ASD relative to TD
Mitotic Nuclear Division
Sister Chromatid Segregation
Nuclear Chromosome Segregation
Mitotic Sister Chromatid Segregation
Spindle Organization
Microtubule Cytoskeleton Organization Involved In Mitosis
Mitotic Spindle Organization
Mitotic Spindle Assembly
Spindle Assembly
Attachment Of Spindle Microtubules To Kinetochore
Regulation Of Neuron Differentiation
Dna Packaging
Hippo Signaling
Centromere Complex Assembly
Regulation Of Microtubule Based Process
Chromatin Remodeling
Neural Precursor Cell Proliferation
Sister Chromatid Cohesion
Chromatin Remodeling At Centromere
Regulation Of Microtubule Cytoskeleton Organization
Protein Dna Complex Subunit Organization
Histone Exchange
Mitochondrial Genome Maintenance
Negative Regulation Of Actin Filament Bundle Assembly
Membrane Docking
Meiotic Cell Cycle
Olfactory Lobe Development
Dna Replication Independent Nucleosome Organization
Glycolytic Process Through Fructose 6 Phosphate
Mitotic Chromosome Condensation
Meiotic Cell Cycle Process
Tube Formation
Positive Regulation Of Arp2 3 Complex Mediated Actin Nucleation
Negative Regulation Of Chromosome Organization
Enriched GO Term
-log10(FDR)
ASD-DM vs ASD-N
ASD-DM vs TD-N
ASD-N vs TD-N

Figure S4

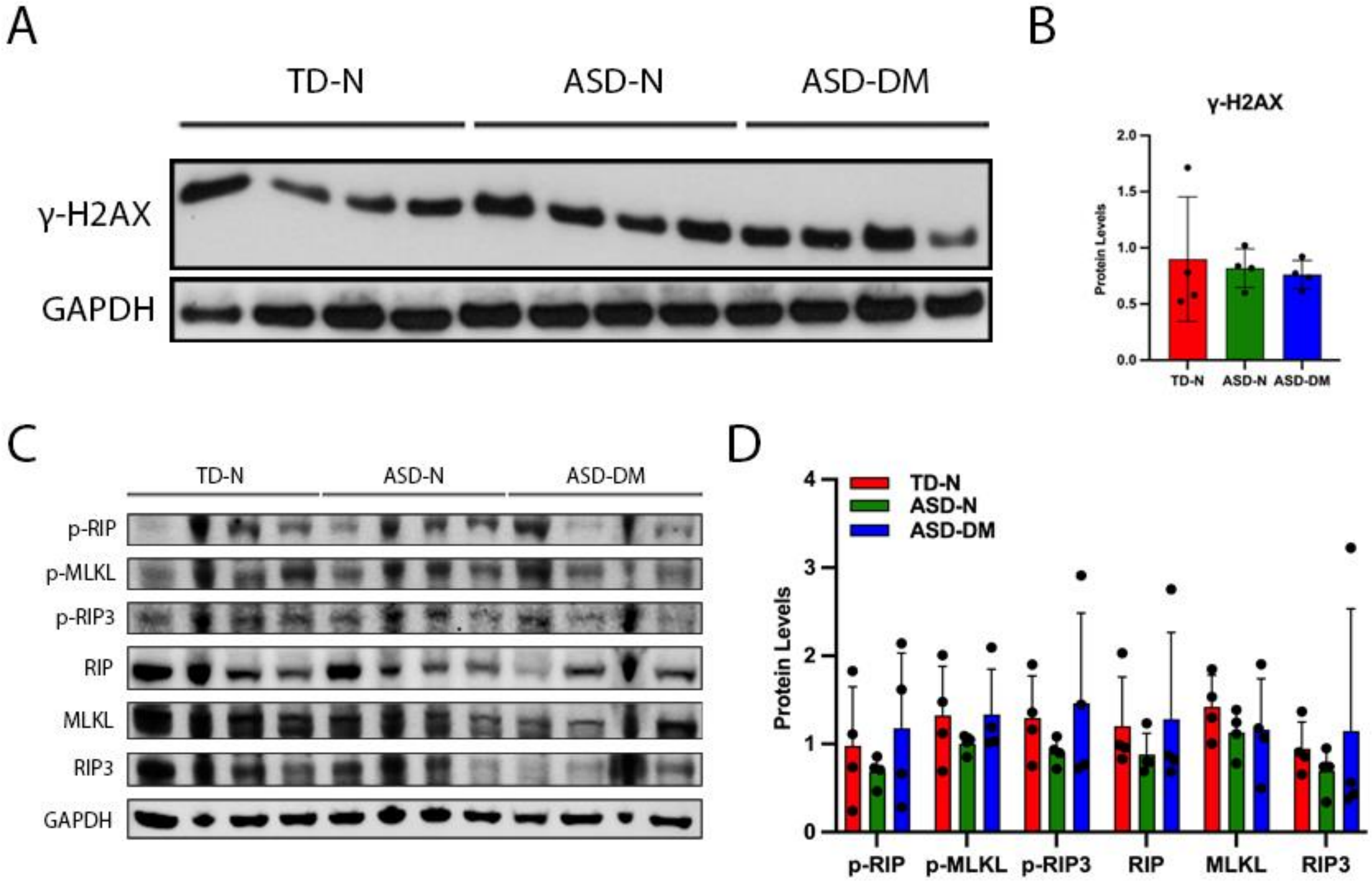

A
TD-N
ASD-N
ASD-DM
γ-H2AX
GAPDH
B
γ-H2AX
Protein Levels
2.0
1.5
1.0
0.5
0.0
TD-N
ASD-N
ASD-DM
C
TD-N
ASD-N
ASD-DM
p-RIP
p-MLKL
p-RIP3
RIP
MLKL
RIP3
GAPDH
D
TD-N
ASD-N
ASD-DM
Protein Levels
4
3
2
1
0
p-RIP
p-MLKL
p-RIP3
RIP
MLKL
RIP3

Figure S5

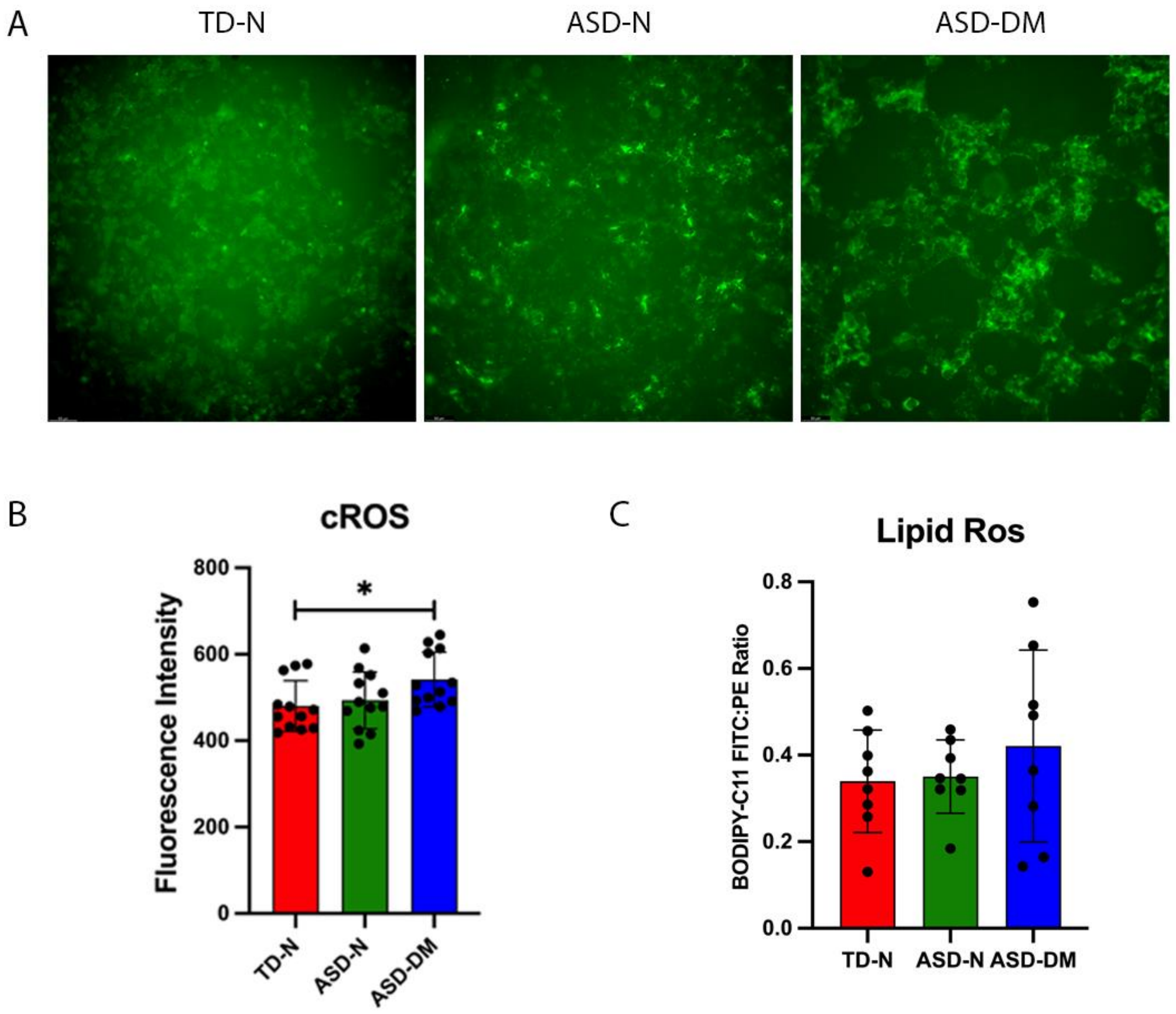
A
TD-N
ASD-N
ASD-DM
B
cROS
Fluorescence Intensity
800
600
400
200
0
*
TD-N
ASD-N
ASD-DM
C
Lipid Ros
BODIPY-C11 FITC:PE Ratio
0.8
0.6
0.4
0.2
0.0
TD-N
ASD-N
ASD-DM

### Figure S6

A

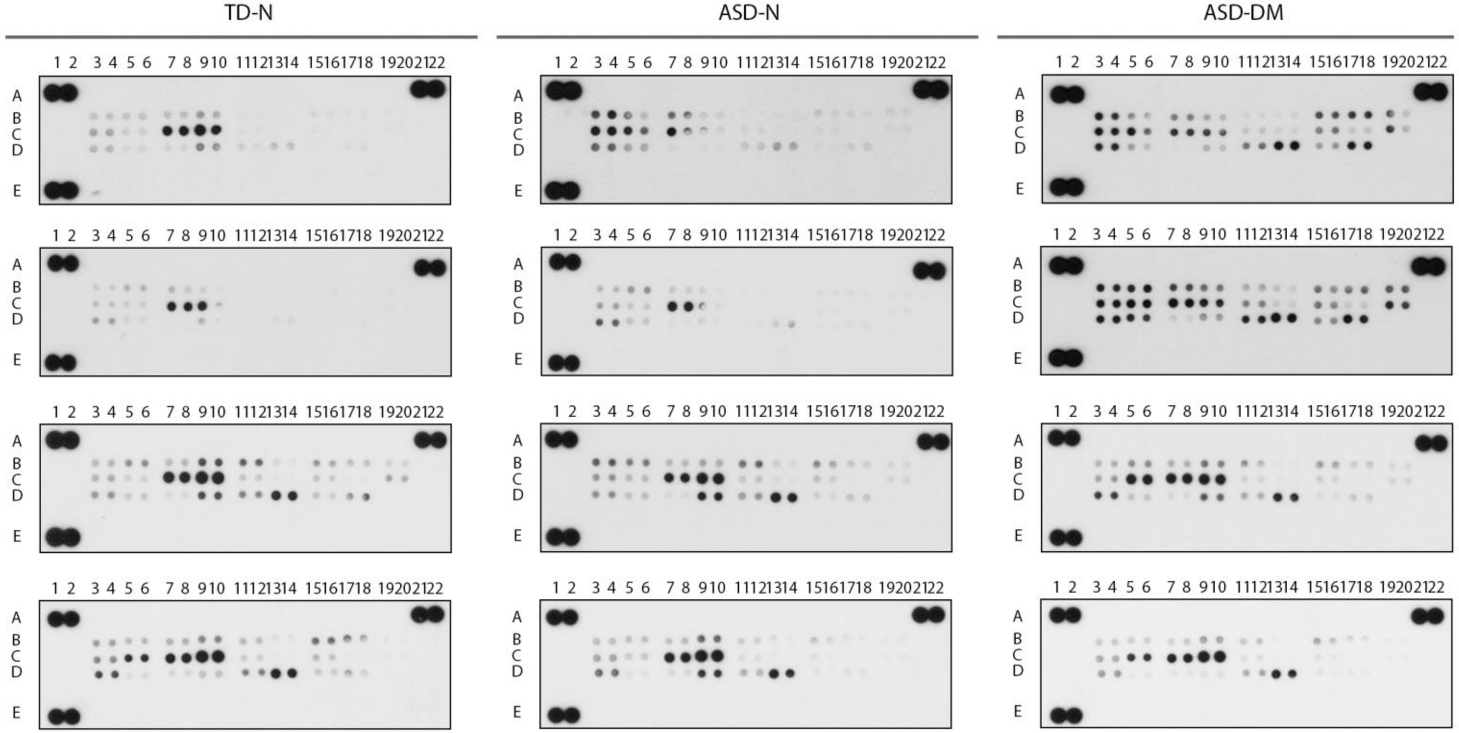


Human Cell Stress Array coordinates

| Coordinate | Analyte/Control |
|---|---|
| A1, A2 | Reference Spots |
| A21, A22 | Reference Spots |
| B3, B4 | ADAMTS1 |
| B5, B6 | Bcl-2 |
| B7, B8 | Carbonic Anhydrase IX |
| B9, B10 | Cited-2 |
| B11, B12 | COX-2 |
| B13, B14 | Cytochrome c |
| B15, B16 | Dkk-4 |
| B17, B18 | FABP-1 |
| B19, B20 | HIF-1α |
| C3, C4 | HIF-2α |
| C5, C6 | Phospho-HSP27 (S78/S82) |
| C7, C8 | HSP60 |
| C9, C10 | HSP70 |
| C11, C12 | IDO |
| C13, C14 | Phospho-JNK Pan (T183/Y185) |
| C15, C16 | NFκB1 |
| C17, C18 | P21/CIP1 |
| C19, C20 | P27 |
| D3, D4 | Phospho-p38α (T180/Y182) |
| D5, D6 | Phospho-p53 (S46) |
| D7, D8 | PON1 |
| D9, D10 | PON2 |
| D11, D12 | PON3 |
| D13, D14 | Thioredoxin-1 |
| D15, D16 | SIRT2 |
| D17, D18 | SOD2 |
| D19, D20 | Negative Control |
| E1, E2 | Reference Spots |

B

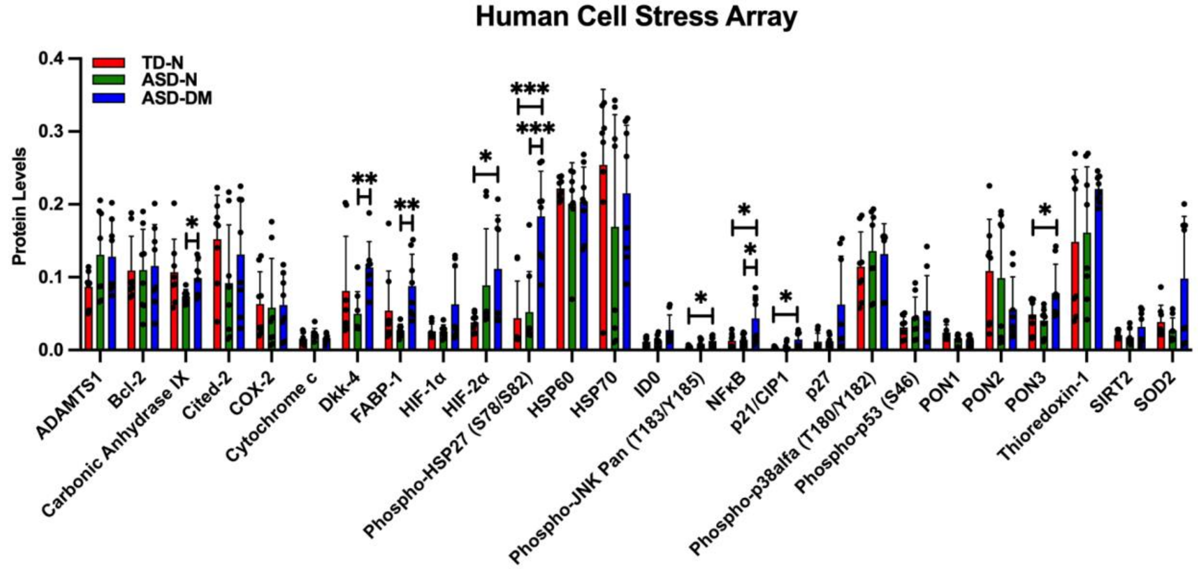

Figure S7

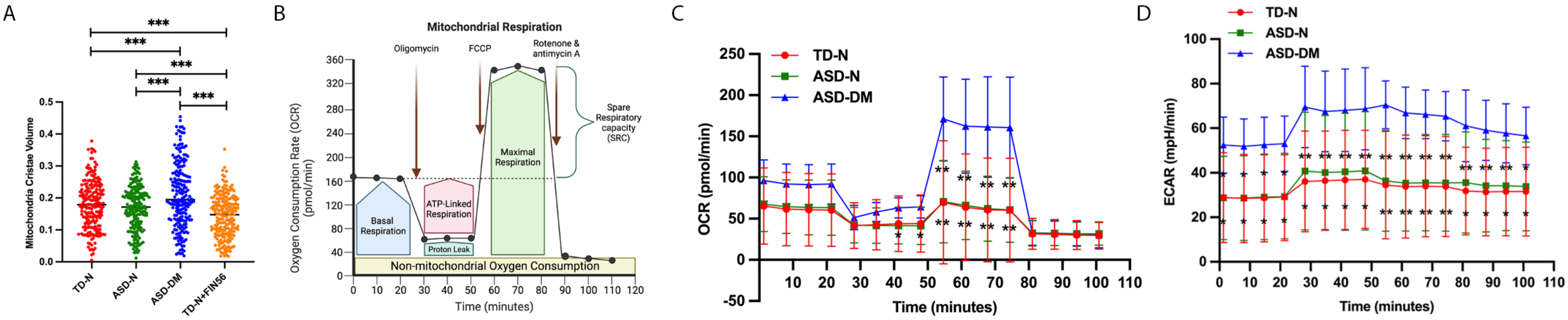
A
B
C
D
***
Mitochondria Cristae Volume
TD-N
ASD-N
ASD-DM
TD-N+FIN56
Mitochondrial Respiration
Oligomycin
FCCP
Rotenone & antimycin A
Spare Respiratory capacity (SRC)
Basal Respiration
ATP-Linked Respiration
Proton Leak
Maximal Respiration
Non-mitochondrial Oxygen Consumption
Oxygen Consumption Rate (OCR) (pmol/min)
Time (minutes)
OCR (pmol/min)
ECAR (mpH/min)

Figure S8

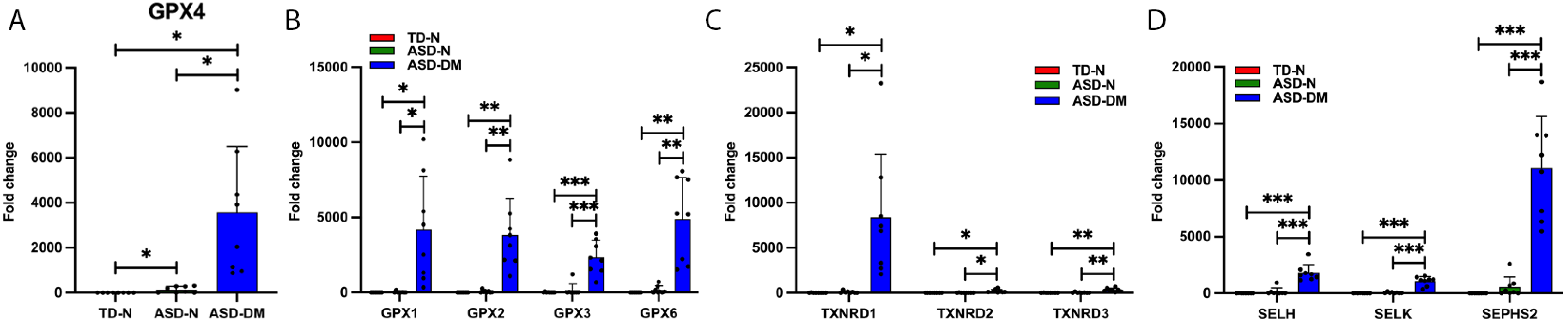
A
GPX4
Fold change
10000
8000
6000
4000
2000
0
TD-N
ASD-N
ASD-DM
B
TD-N
ASD-N
ASD-DM
15000
10000
5000
GPX1
GPX2
GPX3
GPX6
C
25000
20000
15000
10000
5000
TXNRD1
TXNRD2
TXNRD3
D
20000
SELH
SELK
SEPHS2

Figure S9

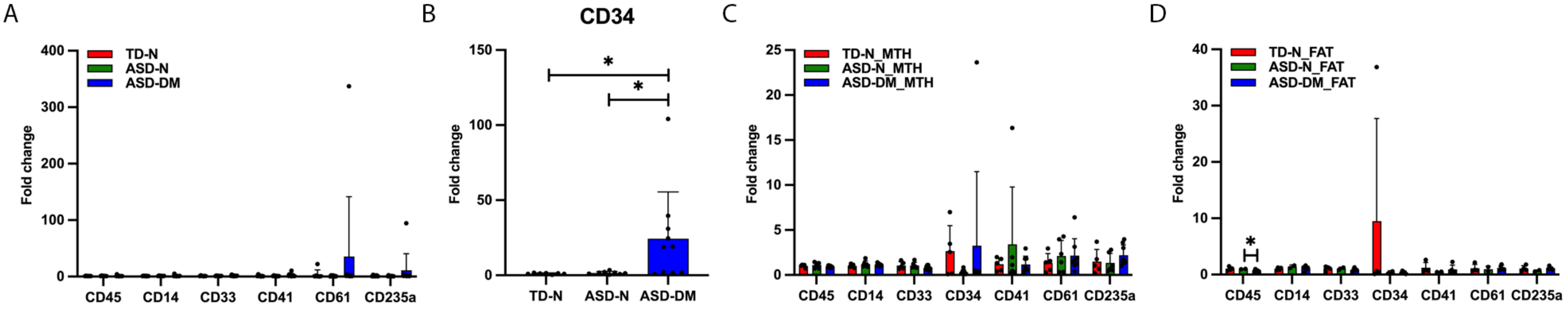

A
B
CD34
C
D
TD-N
ASD-N
ASD-DM
Fold change
CD45
CD14
CD33
CD41
CD61
CD235a
TD-N_MTH
ASD-N_MTH
ASD-DM_MTH
CD34
TD-N_FAT
ASD-N_FAT
ASD-DM_FAT

**Table S1. Clinical characteristics of iPSC subjects**

| Gender | Diagnosis | Subgroup | ADOS Age (months) | ADOS Severity | Age at PBMC (months) | TCV/Height ($cm^3$/inches) |
|---|---|---|---|---|---|---|
| Male | TD† | N | N/A | N/A | 35 | 30.47 |
| Male | TD† | N | N/A | N/A | 30 | 28.30 |
| Male | TD† | N | N/A | N/A | 135 | 30.38 |
| Male | TD† | N | N/A | N/A | 144 | 31.65 |
| Male | ASD | N | 41 | 10 | 42 | 28.77 |
| Male | ASD | N | 39 | 8 | 43 | 30.71 |
| Male | ASD | N | 31 | 10 | 32 | 27.21 |
| Male | ASD | N | 44 | 7 | 139 | 29.46 |
| Male | ASD | DM | 30 | 10 | 35 | 33.63 |
| Male | ASD | DM | 43 | 8 | 43 | 36.11 |
| Male | ASD | DM | 36 | 8 | 121 | 33.71 |
| Male | ASD | DM | 36 | 10 | 117 | 33.69 |

† ADOS was not administered on TD participants.

**Abbreviations**: **TD**: Typical Developing; **ASD**: Autism Spectrum Disorder; **N**: Normal Brain Size; **DM**: Disproportionate Megalencephaly; **ADOS**: Autism Diagnostic Observation Schedule (assessment of communication, social interaction, and play); **TCV/Height**: Total Cerebral Volume ($cm^3$) versus. Height (inches); **N/A**: Not Applicable

**Table S3. Clinical characteristics of whole blood RNA subjects**

| Gender | Diagnosis | Subgroup | ADOS Age (months) | ADOS Severity |
|---|---|---|---|---|
| Male | TD† | N | N/A | N/A |
| Male | TD† | N | N/A | N/A |
| Male | TD† | N | N/A | N/A |
| Male | TD† | N | N/A | N/A |
| Male | TD† | N | N/A | N/A |
| Male | TD† | N | N/A | N/A |
| Male | TD† | N | N/A | N/A |
| Male | ASD | N | 41 | 10 |
| Male | ASD | N | 39 | 8 |
| Male | ASD | N | 44 | 7 |
| Male | ASD | N | 38 | 6 |
| Male | ASD | N | 26 | 9 |
| Male | ASD | N | 36 | 7 |
| Male | ASD | N | 30 | 7 |
| Male | ASD | N | 39 | 5 |
| Male | ASD | DM | 43 | 8 |
| Male | ASD | DM | 36 | 8 |
| Male | ASD | DM | 36 | 10 |
| Male | ASD | DM | 37 | 9 |
| Male | ASD | DM | 38 | 6 |
| Male | ASD | DM | 40 | 6 |
| Male* | ASD | DM | N/A | N/A |
| Male | ASD | DM | 39 | 9 |
| Male | ASD | DM | 39 | 6 |
| Male | ASD | DM | 32 | 7 |

**†** ADOS was not administered on TD participants.

***** Due to masking and the COVID pandemic, this participant was evaluated using the Brief Observation of Symptoms of Autism (BOSA) assessment instead of the ADOS.

**Abbreviations**: **TD**: Typical Developing; **ASD**: Autism Spectrum Disorder; **N**: Normal Brain Size; **DM**: Disproportionate Megalencephaly; **ADOS**: Autism Diagnostic Observation Schedule (assessment of communication, social interaction, and play); **BOSA**: Brief Observation of Symptoms of Autism (BOSA); **N/A**: Not Applicable

**Table S4. Primer sequences for qRT-PCR**

| Gene | Forward Sequence | Reverse Sequence |
|---|---|---|
| *ACTB* | CACCATTGGCAATGAGCGGTTC | AGGTCTTTGCGGATGTCCACGT |
| *CD14* | CTGGAACAGGTGCCTAAAGGAC | GTCCAGTGTCAGGTTATCCACC |
| *CD33* | GTGACTACGGAGAGAACCATCC | GCTGTAACACCAGCTCCTCCAA |
| *CD34* | CCTCAGTGTCTACTGCTGGTCT | GGAATAGCTCTGGTGGCTTGCA |
| *CD41* | CTGTCCAGCTACTGGTGCAAGA | ATGTTGTGCCCAGTGGCTCCAA |
| *CD45* | CTTCAGTGGTCCCATTGTGGTG | CCACTTTGTTCTCGGCTTCCAG |
| *CD61* | CATGGATTCCAGCAATGTCCTCC | TTGAGGCAGGTGGCATTGAAGG |
| *CD235a* | ATATGCAGCCACTCCTAGAGCTC | CTGGTTCAGAGAAATGATGGGCA |
| *GPX1* | CAGTCGGTGTATGCCTTCTCG | GAGGGACGCCACATTCTCG |
| *GPX2* | GAATGGGCAGAACGAGCATC | CCGGCCCTATGAGGAACTTC |
| *GPX3* | GAGCTTGCACCATTCGGTCT | GGGTAGGAAGGATCTCTGAGTTC |
| *GPX4* | GAGGCAAGACCGAAGTAAACTAC | CCGAACTGGTTACACGGGAA |
| *GPX6* | GCCCTCCGACCTCTGATCTT | TGTCTGACTTGACTGTGCTGA |
| *SELH* | GCTTCCAGTAAAGGTGAACCCG | ACCCAAATCTCCCTACGACAGG |
| *SELK* | TCTGGGGAATAGCTGAGTTTGT | CACGCAGATGATTGATTCTACCC |
| *SEPHS2* | GCAGACCACGGACTTCTTTTA | ACTCAGCACGTTGGCACAA |
| *SOX1* | CAGTACAGCCCCATCTCCAAC | GCGGGCAAGTACATGCTGA |
| *TXNRD1* | ATGGGCAATTTATTGGTCCTCAC | CCCAAGTAACGTGGTCTTTCAC |
| *TXNRD2* | GTGGGCCATAGGTCGAGTC | TGAGTGTCGGGGCTAGTATCTA |
| *TXNRD3* | ATGACAAAAGCGATTCAGAACCA | ACCCGTTGCTATGACAAACTG |
